# Advanced Ellis Concept for a Fiber-Optic Fluorescent Microscope

**DOI:** 10.64898/2026.04.10.717647

**Authors:** Aleksey Klepukov

**Author notes:** corresponding author: DiI-lab (private company). Head of private company. Russia, Moscow, Kropotkinskiy pereylok, 20; 119034.

## Abstract

The design of the classical fluorescence microscope has undergone few changes since the 1970s-1980s, when Ploemopak modules with filter cubes became widespread. Most of these changes have been in the replacement of mercury and xenon lamps with LED illuminators in the 2010s. However, this does not mean that this stable design cannot be improved upon. New method: The implementation of a vibrating optical fiber, positioned using a micromanipulator and connected to any suitable type of laser, enables a full spectrum of fluorescence research. This work presents an advanced version of the Ellis concept, in which light is delivered directly onto the sample, rather than into the filter cube (technical novelty).To confirm the functionality of the microscope, vibrational slices of mouse brain stained with three fluorescent markers (B3-PPC, DiI and DiD) covering most of the visible spectrum were examined. The fiber-optic illumination system eliminates the need for bulky and obsolete high-voltage plasma arc lamp units without compromising image quality (confirmed by the USAF 1951 test and SDNR assessment on fluorescent beads). Furthermore, the optical fiber mounted on manipulators is convenient and easy to integrate, for example, into stereomicroscopes for scanning large brain tissue samples.

## 1. Introduction

The main trends in modern fluorescence microscopy lie in the field of high and super-resolution at the boundary of or beyond the diffraction limit. Confocal microscopy, the era of which began in the second half of the 1980s (the first major review publication on this topic was released in 1990 – Handbook of Confocal Microscopy, First edition), gave a tremendous technological impetus to the entire field of fluorescence microscopy. However, this impetus did not affect the basic concept of the classical fluorescence microscope, which stabilized at the level of the 1970s (when filter cube modules with fluorescence cubes became widespread). The only very relative improvement in the design of the basic fluorescence microscope concept occurred in the 2010s - this was the replacement of mercury and xenon lamps with LED illuminators. The rest of the filter cube module remained unchanged. Does this mean that the current concept of the fluorescence microscope is ideal and that nothing from confocal microscopy can be incorporated into it? And most importantly – why incorporate anything, if the confocal microscope is itself the logical next step in the development of the fluorescence microscope?

Let’s start with the fact that the fluorescence microscope is still actively used in routine research, i.e., this technology is relevant and meaningful. There are several problems with the concept of the classical fluorescence microscope, and all of them stem from the age of this technology (a half-century of minimal changes is taking its toll).

a. Accessibility of the method. The main problem. It is impossible to simply convert any transmitted light microscope into a fluorescent one, because a filter cube block is not designed for every microscope model. Furthermore, filter cube blocks cannot always be installed even on different models from the same manufacturer (e.g., due to different mounting diameters). So, if a manufacturer has not developed a filter cube block for a specific microscope model, that model is essentially excluded from fluorescence imaging, even if it is excellent in all other respects. This is especially true for stereomicroscope models, for the overwhelming majority of which filter cube blocks are not developed at all.
b. Spectral limitations. The filter cubes produced for a particular microscope model impose rigid limitations on visualizing the spectrum of the molecules under study. While cubes for 400-550nm (in the excitation spectrum) are quite common, the same cannot be said for cubes in the 590-650nm range. This work, in fact, originated as a reaction to the author’s inability to quickly find a 650nm filter cube for evaluating the fluorescent marker DiD (see results with this marker later) due to its absence from product lines.
c. Power supply limitations. A mercury lamp requires a high-voltage power supply with a lamp ignition system. Such power supplies are still expensive, unavailable for general purchase outside specialized microscopy equipment manufacturers, and often cannot be ordered separately from the lamphouse. LED illuminators are less demanding in this sense, but they are still significantly less common and have not been developed for all models. This article also arose, in part, as a reaction to the inability to quickly replace a burned-out power supply for a Leitz Labovert with a newer one without modification (to the cable connector), which was also difficult to obtain.
d. The mercury lamp also has its drawbacks. Firstly, due to its high cost (usually over $100), and secondly, due to its very limited service life (200 hours for a 100W lamp like Osram or 100 hours for a DRSH-100-2; service life information is taken from official manuals). Furthermore, the plasma ball in the center of the lamp is usually unstable in position; it always fluctuates slightly, creating some inconvenience when evaluating a sample due to inevitable changes in object illumination brightness (Handbook of Confocal Microscopy, 2006). Additionally, if the lamp’s seal is compromised, there is a risk of extremely toxic mercury vapor evaporation, which can harm the operator’s health.

Thus, the problems are clearly outlined, there are many of them, and solutions must be sought. And such a solution was found. Even in the very first edition of the Handbook of Confocal Microscopy (1990), the fundamental schematic by Ellis (1979) was presented, showing the delivery of laser light through a vibrating multimode optical fiber to the objective (via a dichroic mirror). For unknown reasons, this concept is rarely mentioned in the literature, although isolated studies do exist (Reitz and Paglario, 1994; Silvrmintz et al., 2003). This work aims to vividly highlight the advantages of the Ellis concept, especially in applications to objects in neuroscience.

The novelty of the method:

1. Incorporating positional micromanipulators into the design (partially implemented by Nightsea, but in an simplified version and only for stereomicroscopes - https://nightsea.com; Masel 2013). Micromanipulators allow for exceptionally precise alignment of the illumination area onto any region of interest and, for example, enable illuminating the object with the very edge of the bright field (see results), gradually shifting the position of the bright field.
2. Because the illumination field can be easily and arbitrarily controlled, it becomes possible to scan the illuminated area across the sample (including non-standard sized ones) in search of labeled regions (see results).
3. The manipulators also allow for the elimination of filter cubes (since the optical fiber can be directed straight at the object), requiring only emission filters. Consequently, non-standard combinations of lasers and emission filters become possible, which are unavailable in the standard set of cubes for a fluorescence filter block (restrictions imposed by the dichroic mirror are removed, see results for the USAF 1951 test).

The introduction of optical fiber and micromanipulators removes limitations both in terms of spectrum and method accessibility (micromanipulators can be mounted on any microscope using an aluminum structural framework), as well as complexities related to power supply. The issue with toxic mercury vapors also disappears on its own, as there is no mercury lamp.

Vibratome slices of mouse brain (100 µm thick), stained with various fluorescent markers covering a large portion of the visible spectrum—namely B3-PPC, DiI, and DiD—were used as the subject for comparing the functionality of classical and fiber-optic fluorescence microscopes. Furthermore, to demonstrate the capability of scanning a sample using micromanipulators, a large slice of calf brain was employed.

To prove the advantages of the Ellis concept compared to traditional fluorescence microscopy, a series of tests on non-biological objects was additionally conducted, specifically:

1. Measuring fluorescence intensity using a photometric attachment (ФМЭЛ-1A) to obtain precise quantitative data on the differences between illumination systems.
2. Imaging fluorescent beads with diameters of 200 nm, 3 µm, and 10 µm to determine the limits of optical resolution and identify the signal-to-noise ratio.
3. An optical resolution test using the USAF 1951 target.

## 2. Materials and methods

### 2.1. Bioethics commission

In this study, the Bioethics commission of the Faculty of Biology of Moscow State University has approved all procedures on animals. Such procedures on animals as perfusion and euthanasia were carried in the Institute of the Mitengineering of the Moscow State University in accordance with the Directive 2010/63 / EU of the European Parliament and the Council of the European Union for the protection of animals used for scientific purposes. All the work was carried out on mice - hybrids F1 - C57Bl6/CBA. The total number of animals was 7. The birthday of the mice was considered a one postnatal day P1, all procedures were performed on mice on the eleventh postnatal day P11. A list of all mice used and manipulations performed on them is given in Table 1.

**Table 1:**
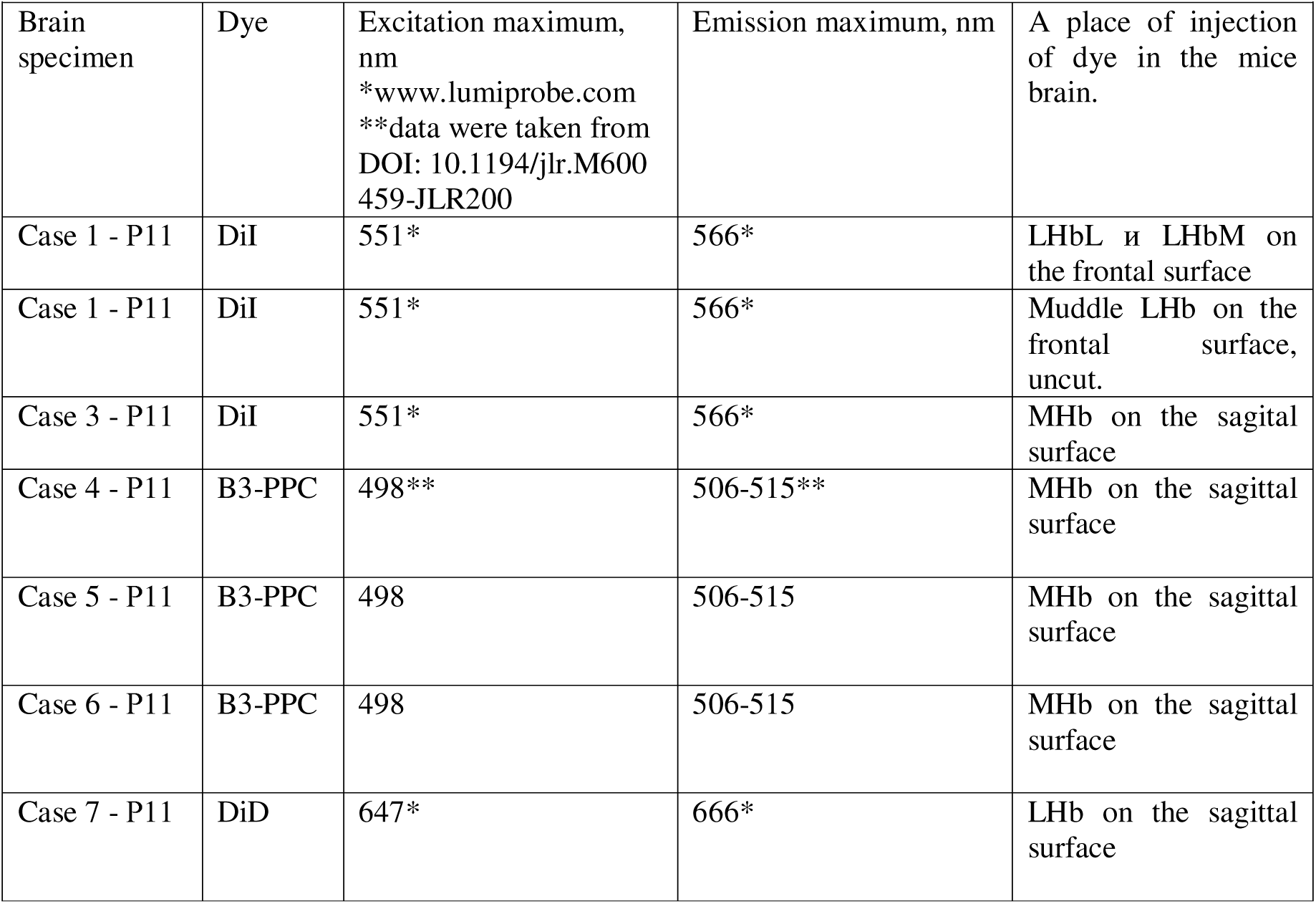
Mouse brain samples used in this study depending on the research aim of the study.

|  |  |  |  |  |
| --- | --- | --- | --- | --- |
| Brain specimen | Dye | Excitation maximum, nm<br>*www.lumiprobe.com<br>**data were taken from<br>DOI: 10.1194/jlr.M600459-JLR200 | Emission maximum, nm | A place of injection of dye in the mice brain. |
| Case 1 - P11 | DiI | 551* | 566* | LHbL и LHbM on the frontal surface |
| Case 1 - P11 | DiI | 551* | 566* | Muddle LHb on the frontal surface, uncut. |
| Case 3 - P11 | DiI | 551* | 566* | MHb on the sagittal surface |
| Case 4 - P11 | B3-PPC | 498** | 506-515** | MHb on the sagittal surface |
| Case 5 - P11 | B3-PPC | 498 | 506-515 | MHb on the sagittal surface |
| Case 6 - P11 | B3-PPC | 498 | 506-515 | MHb on the sagittal surface |
| Case 7 - P11 | DiD | 647* | 666* | LHb on the sagittal surface |

In addition, one sample of calf brain were used (one sagittal half of the whole brain cut along the midline). The sample was purchased at the commercial market in Moscow (Danilovsky Rynok, JSC, TIN 7725582006) in accordance with all regulatory procedures for biological material transfer.

### 2.2. Procedures performed on mice

All mice (P11) were given intraperitoneal solutions of zoletil 100 (Vibrac) in saline (0.9% NaCl) at the rate of 10 mg per 1 kg of body weight before perfusion. Then mice were subsequently perfused with saline solution (0.9% NaCl) through the left ventricle of the heart to remove blood from the blood vessels. Thereafter, the mice were decapitated and the heads were placed in 4% paraformaldehyde (PAF, PanReac AppliChem, СAS 30525-89-4) on 0.1 M PB (pH 7.4) for up to one month term at room temperature for postfixation. After postfixation, the brain was removed from the skull and stored in 4% PAF.

### 2.3. Histological part – Mice brain

1. For frontal marker applications, a mouse brain sample was placed in a warm (45°C) solution of 5% low-melting agarose (Serva CAS [9012-36-6]). After the agarose solidified, a cubic block with the brain was cut out of it, which was then attached to the bottom of the 3D-vibrting microtome tub using ordinary superglue (its design will be described later). The tub was filled with saline solution (0.9% NaCl) after the glue solidified. Then the block was cut on the 3D-vibrating microtome in a rostrocaudate direction with a trapezoidal blade (Dexter SK5, blade angle 25-30 degrees) to the desired level (corresponding to the Habenula level).
2. In the case of sagittal marker applications, the mouse brain sample was cut manually along the midline with a trapezoidal blade.
3. After the desired brain level was found, the bath with the agarose block was removed from the 3D-vibrating microtome and the saline solution was drained. The block with the brain was cut and after the cut surface dried, a lipid fluorescent marker was applied to the desired area. The markers were applied with a glass needle (SOOK Medical) installed in a Narishige MN188 positioning micromanipulator. All applications were performed under the control of a stereomicroscope (AmScope 07-40).
4. Overall, three different lipid fluorescent markers were used for the applications - B3-PPC (1-palmitoyl-2-[(Me4-BODIPY-8)-acyl]-sn-glycero-3-phosphocholine with acyl containing 3 carbons, the preparation was synthesised and provided by the Institute of Bioorganic Chemistry of the Russian Academy of Sciences). DiI 1,1′-Dioctadecyl3,3,3′,3′,3′-Tetramethylindocarbocyanine Perchlorat, Lumiprobe, Cas number 22366-93-4); DiD (Lumiprobe, Cas number 75539-51-4). DiI and DiD were applied in dry crystalline form, B3-PPC was applied as a viscous oily liquid (in this case, immediately after application, the brain block was reconnected in 5% agarose). All applications were made in Hb, the list of applications is given in Table 1.
5. Brain agarose blocks with marker applications were stored in 4% PAF for 0.1M PB for a period of 2 months at a temperature of 25°C. After the incubation period required for diffusion of the markers along the membranes, the agarose blocks with brains were extracted from formaldehyde and sliced on a 3D-vibrating microtome into frontal or sagittal slices of 100 μm thickness (Case 6 sliced into sagittal slices of 50 μm thickness).
6. All mouse brain sections were placed under a coverslip in mounting medium (Mowiol-4.88 mounting medium; Kremer; CAS number 25213-24-5. The 2.5% glycerol-based Mowiol-4.88 solution is a standard (http://cshprotocols.cshlp.org/content/2006/1/pdb.rec10255). If one or two manipulators were intended to be used, standard slides were used.

### 2.4. Histological part – Calf brain

A fresh brain sample (one sagittally cut half) was cut with an ordinary kitchen knife into fragments approximately 82 cm (96 cm3 volume); these fragments were then immersed in 4% PAF for at least one week. Prior to cutting, the cut brain fragment was dissected, using tweezers, from the dura mater, primarily from the cerebral vasculature. After removal of the vascular membrane, excised brain fragments 96 cm3 volume were poured into a fusible agarose block with a final size of 94 cm (252 cm3 volume). After the block was attached to the bottom of the bath of the 3D-vibrating microtome with superglue, the frontal surface was oriented to the bottom, the dorso-ventral axis was perpendicular to the cutting surface of the blade. A segmented construction knife blade, Dexter SK5 (blade width 25 mm, cutting surface length 120 mm), was used to slice the selected calf brain slice into slices of 150 μm thickness. The angle of inclination of the blade was 25°.

After slicing the brain tissue block, small DiI crystals were applied with a steel needle from an insulin syringe to various points on the surface of a randomly selected calf brain slice (at the level of the rostral part of the capsula interna). Immediately after, technical acetone (ГОСТ 2768-84, 10 µl per crystal) was applied to these crystals to dissolve them within the thickness of the slice. Technical acetone destroys cell membranes, therefore, for revealing fine cellular structures on the slice, it is not only useless but harmful. However, in this case, the goal was precisely rapid and durable staining of random areas on the slice surface to demonstrate the scanning capabilities of the fiber-optic microscope. Within five minutes of applying the acetone droplets, the slice was rinsed under running water and, without drying, mounted using a mounting medium (Mowiol-4.88 mounting medium; Kremer; CAS number 25213-24-5. The 2.5% glycerol-based Mowiol-4.88 solution is a standard (http://cshprotocols.cshlp.org/content/2006/1/pdb.rec10255) between two 100×100mm round coverslips.

### 2.5. Specimen analysis

Evaluation in all cases was performed through an Leitz PL objective (2.5x); Leitz EF objectives (4x) and Zeiss Neofluar objective (10x), China LM Plan objective (20x), China LM Plan objective (50x), Leitz Fluotar objective (100x).

Digital images of microscope slides of a mouse brain were obtained using a MG3CMOS06300KpA-2018 camera (ISOLAB 6.3MP, Sony IMX178 (C) sensor) and ImageView software.

In addition, an Allied Vision Marlin F145B2 digital machine vision camera (1.4MP, Sony ICX205 (B) sensor) with proprietary Vimba View software was used.

Computer program Photoshop CS3 (Adobe, USA) was used for processing and editing of digital images and for illustration of the results.

### 2.6. Description of the design of a fiber optic fluorescence microscope (advanced Ellis concept)

The concept of any modern fluorescence microscope is based on an optical scheme where light from a mercury lamp (or an LED illuminator) is transmitted through a focusing collimator (and a neutral density filter) onto a dichroic mirror. From the dichroic mirror, the light is then focused through the objective lens onto the specimen (Figure 1.A). The idea of Ellis (1979) is to direct a concentrated light source from a laser (instead of from a mercury lamp) onto the dichroic mirror of the filter cube via a vibrating optical fiber (Fig.1.B), bypassing the system of lenses in the collimator and the neutral density filter, which become unnecessary since the laser provides a bright, monochromatic, and narrow-band beam of light. The vibration of the optical fiber serves to mitigate the phenomenon of laser speckle. The concept of the fiber-optic fluorescence microscope is a combination of these two schemes (Fig.1.C).

**Figure 1.**
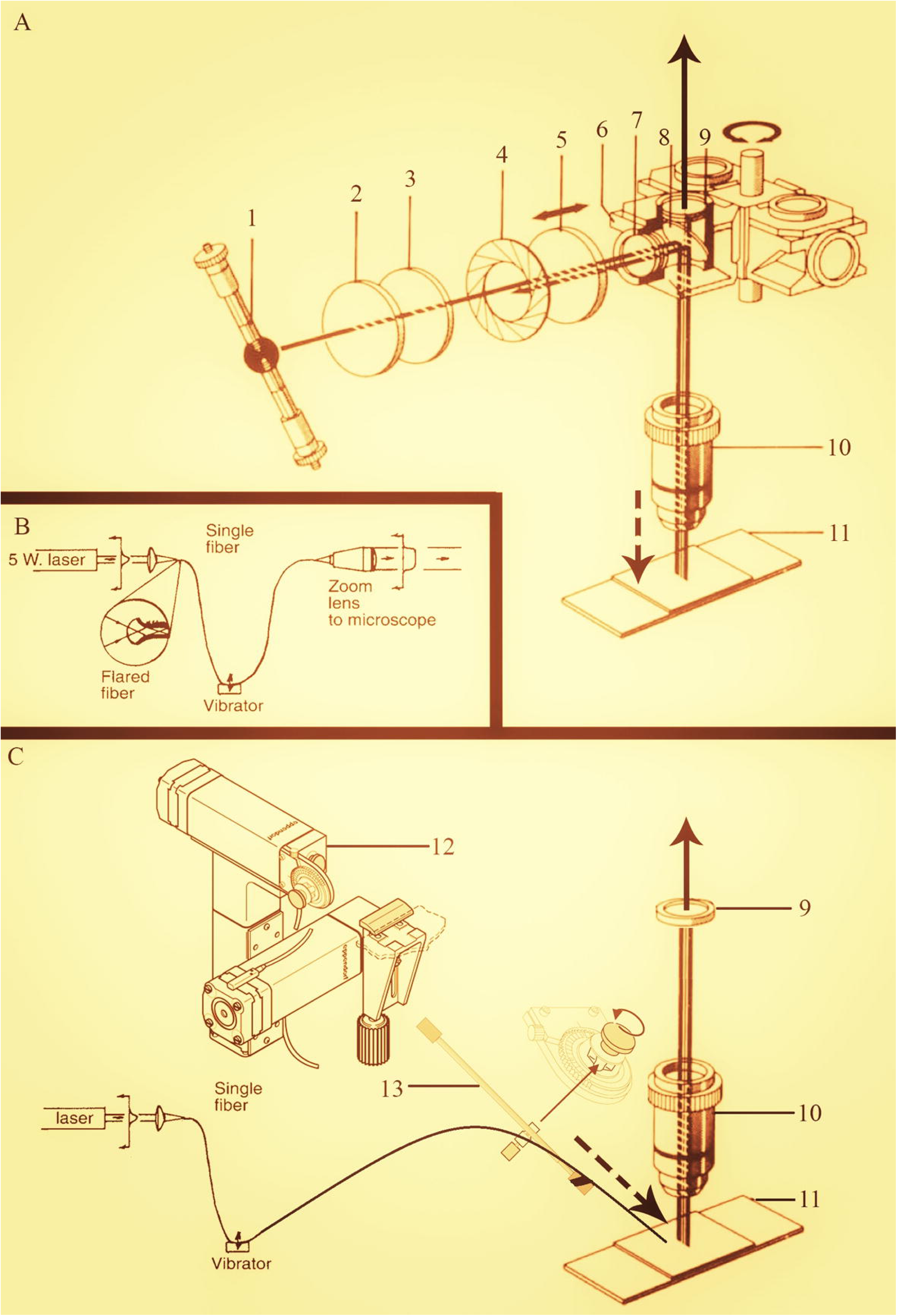
A – A classical optical scheme with epifluorescent illumination of the specimen through the objective lens using a mercury lamp, a collimator, and filter cubes. The numbers in the figure denote system components: 1 – light source; 2 – heat filter; 3 – Red suppression filter; 4 – field diaphragm; 5 – lens; 6 – filter cube; 7 – exiting filter; 8 - dichroic mirror; 9 – suppression filter; 10 - objective; 11 – specimen. Image taken from the official instruction manual for the Leitz Diaulux microscope. B – The original optical scheme by Ellis (1979) with illumination of the specimen via an optical fiber connected at one end to a laser and with its output directed into the microscope optics by the other, free end. The fiber itself is connected in the middle to a vibrational module. The scheme is taken from the Handbook of Confocal Microscopy (2006). C – An optical scheme of a fiber-optic fluorescence microscope that combines the Ellis concept with the classical epifluorescence microscope scheme, ultimately requiring radically fewer components (denoted by numbers); in the limiting case, only an emission filter remains. A micromanipulator (12, Eppendorf model) is added, which positions the free tip of the optical fiber (mounted on a holder 13) directly near the specimen, from the side. The fiber itself is single-mode. The design allows for the positioning of up to 6 fiber-optic tips around the specimen. The dotted bold arrow is the exciting light; the continuous bold arrow is emission light.

In a fiber-optic fluorescence microscope, low-power (300-1000 mW) laser pointers or laser modules based on laser diodes of different spectra (blue at 450-490 nm, green at 532 nm, or red at 650 nm) with a collimator already built into the pointer are used instead of a mercury lamp. Their advantages are simple design and low cost. A single-mode optical fiber is connected to the laser pointer, and its other (free) end is positioned using a micromanipulator directly near the object (Figure 1.C). This design eliminates the need for a focusing collimator, an excitation filter, and a dichroic mirror in the filter cube (and indeed the filter cube itself, with only an emission filter being required). In this setup, the microscope objective is not relevant as a light-focusing device because the optical fiber is located very close to the object (to the side of it), and light loss due to the scattering effect is minimal. The absence of filters and lenses in the fiber-optic path (the collimator of the laser pointer does not count) also leads to minimal loss in radiation intensity (whereas in a standard fluorescence microscope setup, the light from a mercury lamp is attenuated approximately a million times by the time it exits the objective; information taken from the Handbook of Confocal Microscopy, 2006). A vibration module is retained to mitigate the laser speckle effect.

In summary, the fiber-optic fluorescence microscope can be broken down into a number of modules (Fig.2.А). The module design is very simple and does not require a large number of components, which are often supplied pre-assembled. All that is needed is to connect the modules in the correct sequence.

1. Module 1 - Laser Fiber Unit. As a light source in the fiber optic fluorescence microscope are used ordinary household laser pointers of low power (300-1000mW), with a wavelength of 450nm (blue laser), 488nm (turquoise laser), 532nm (green laser) and 650nm (red laser). A single-mode optical fiber (thickness 2mm, length 1 mester) is attached to the laser pointer through an adapter (a fitting with a twisted skein of tape inside), the other free end of the optical fiber is attached to the end of the metal rod of the micromanipulator with tape.
2. Module 2 – Micromanipulators. Three mutually perpendicular linear translators SEMX-60AC were taken as a basis for the micromanipulator (on the y-axis macro-rails can be used). The z-axis translator is screwed to the z-axis translator with a metal rod, to the end of which the free tip of the optical fiber is glued with ordinary duct tape. One can also use any ready-made commercial micromanipulators, for example from Eppendorf or Narishige.
3. Module 3 - Vibration Unit. A vibrator creating low-frequency vibrations (10-20Hz) is attached to the single-mode optical fiber at any convenient place along its course. The vibrator can be a design based on the crank mechanism of a 3D vibration slicer, connected to a separate power supply unit. It is also possible to use as an alternative any ready-made linear motion module with a ZK-SMC02 programmer in oscillation mode.

**Figure 2.**
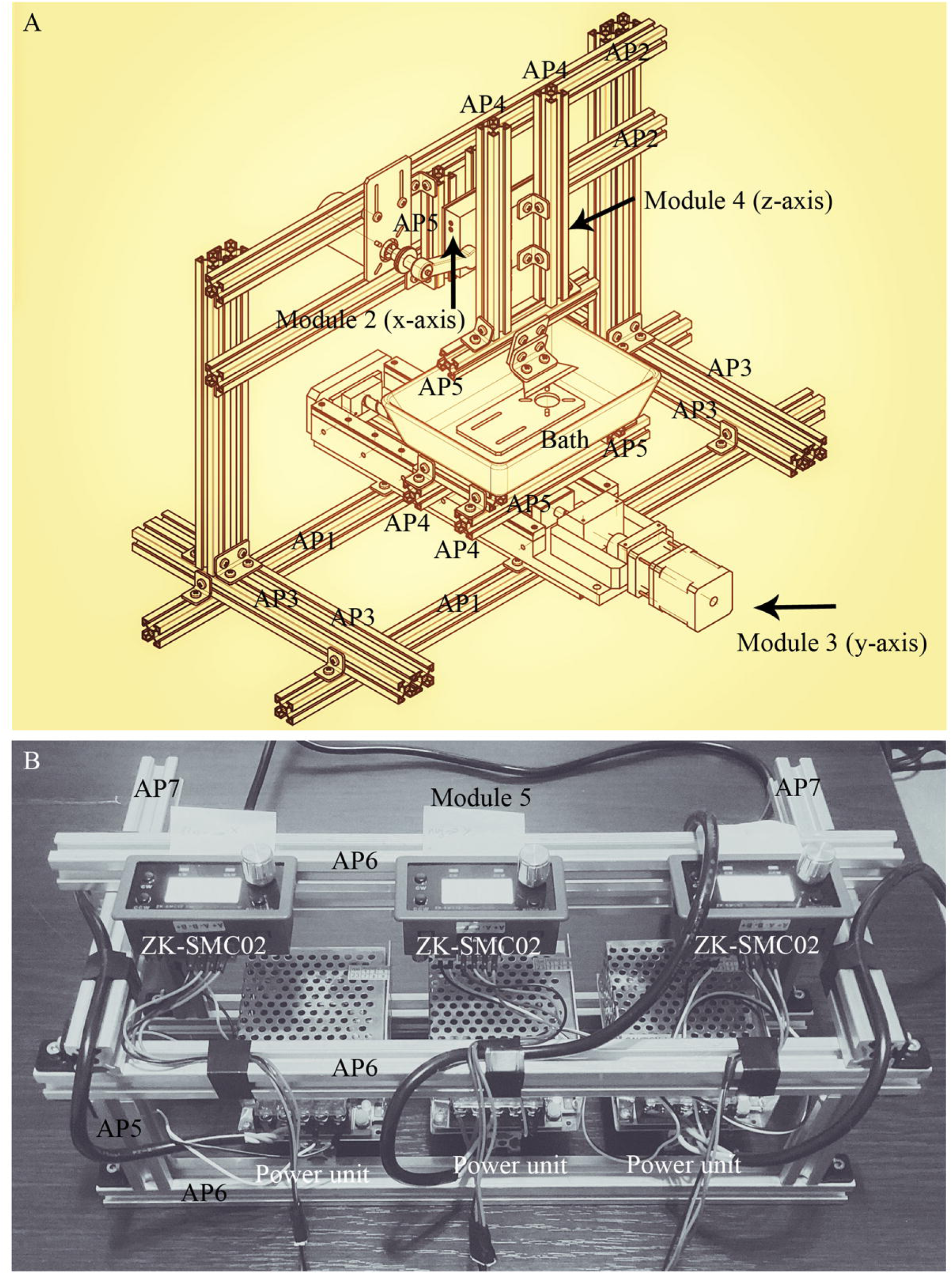
A - general view of a fiber-optic fluorescence microscope, based on the Leitz Laborlux S model (Module 1) with two micromanipulators mounted in the upper position. The micromanipulators are attached to the microscope slide via an aluminium metal construction adapter, and provide a single-mode optical fiber from the lasers as close and precise as possible to the object. The vibration unit is not shown in the frame. The emission filter is located in the housing from the Ploemopak unit. Detailed description of the parts is given in Table 2. B – An independent, general view image (separate from the microscope) of a universal kit for converting a transmitted-light microscope into a fiber-optic microscope. The kit includes: a - a set of lasers (model chosen by the laboratory); b - single-mode optical fiber branched at one end (or unbranched, resulting in a one-laser-one-fiber setup); c - a vibration module for the optical fiber; d - electronic manipulators (Eppendorf 5171 or any suitable linear translation stages with stepper motors). The photo does not show interference filters (they are typically located in the optical path within the microscope body) and the laser power supply. The control system for the electronic manipulators (any type) is shown in Figure 4.B. The kit in the photo is sufficient to convert any transmitted-light microscope into a fiber-optic microscope.

**Table 2.**
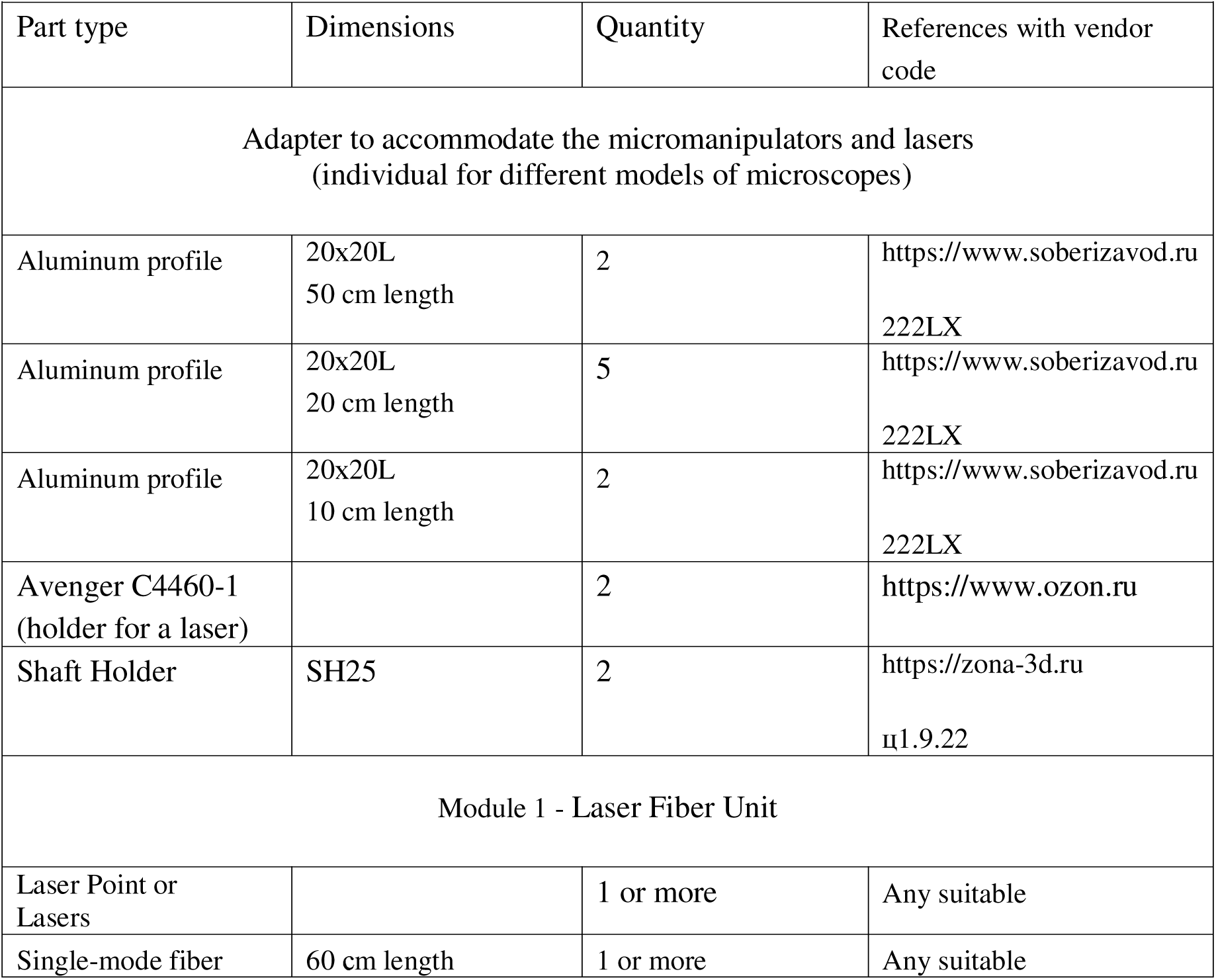
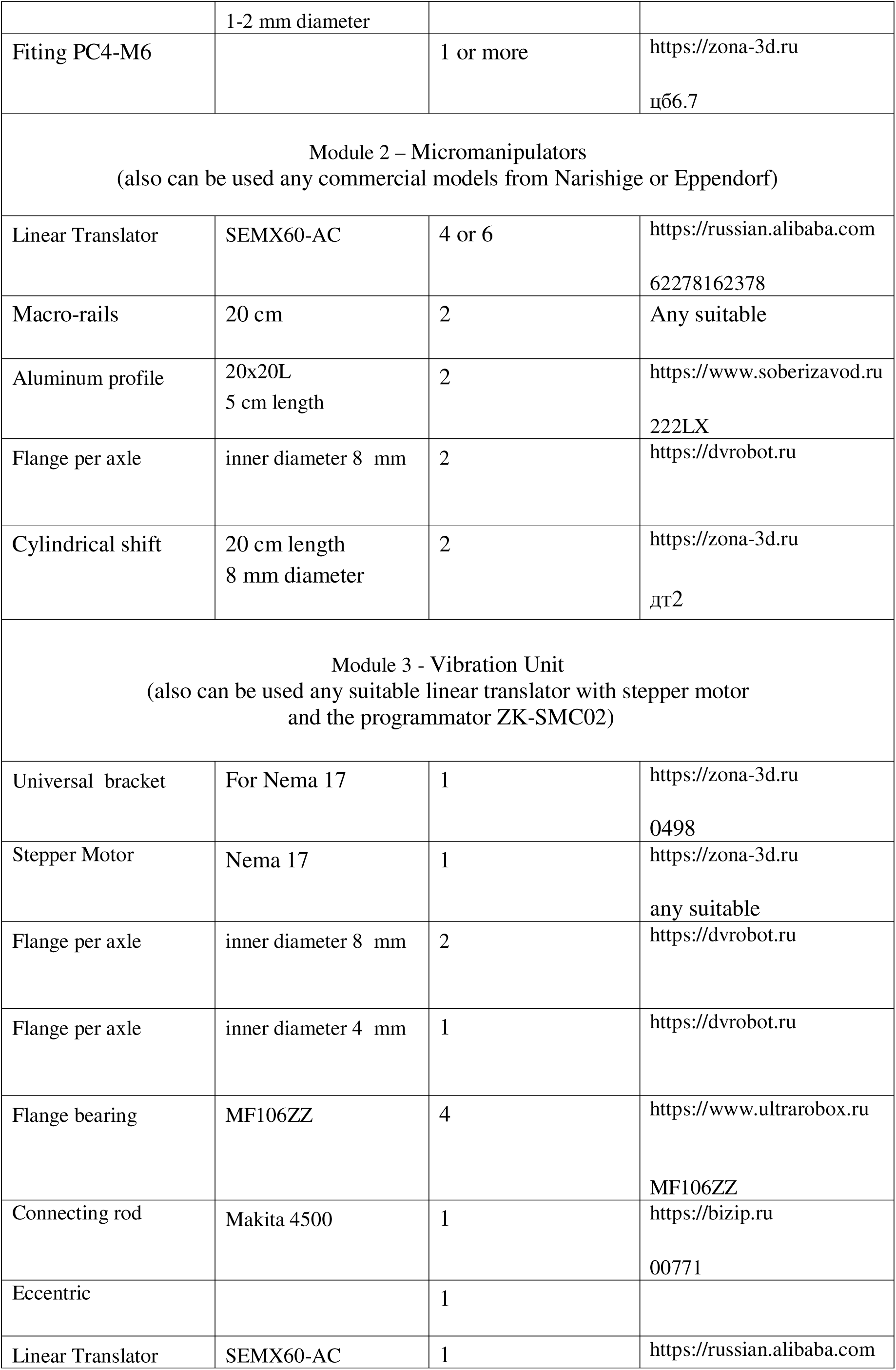
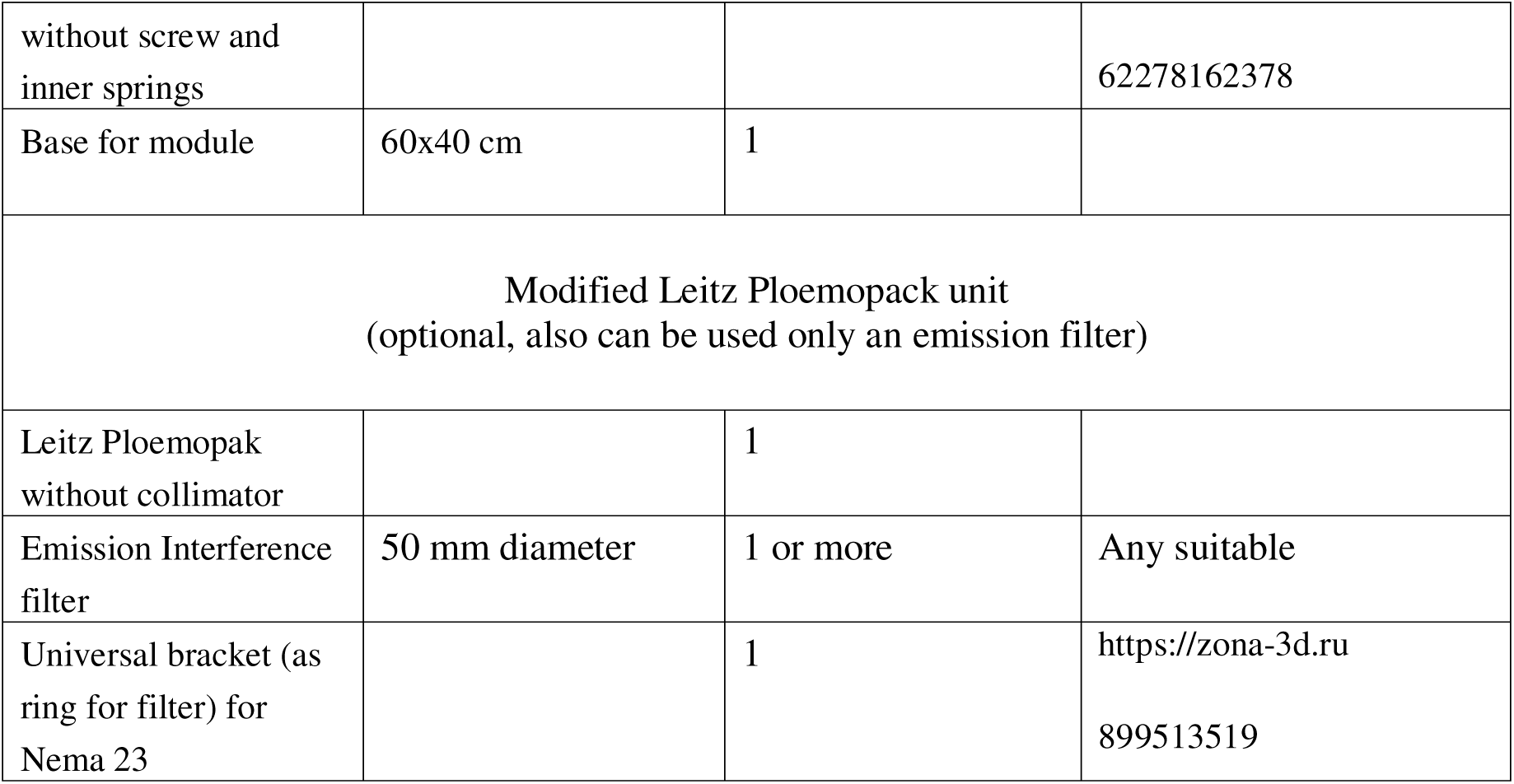
List of necessary parts of the corresponding abbreviations required to build an Optic-fiber fluorescent microscope. The list of parts is divided into modules that form the microscope.

Thus, all three modules are supplied either ready-made (the vibration module on a linear translator and micromanipulators) or are very easy to assemble (by connecting the optical fiber to the laser). All three modules are mounted onto the microscope body using an aluminum frame kit (the number of frame components and their orientation depend on the microscope model and are selected individually).

It should be separately noted that the design of the fiber-optic fluorescence microscope allows for either preserving the original Ploemopak module with cubes (where only the emission filter in the cubes is utilized) or its complete replacement, since the emission filters can be installed directly into the objective of the digital camera (which is used to capture the object). In this work, an intermediate option was chosen: all the contents were removed from the Ploemopak module and the collimator was detached. I.e. only the body and the optical system of two lenses on the way from the lens to the eyepieces remained from the Ploemopak. A ring was placed between the lenses on which an emission interference filter was placed. A total of three Carl Zeiss interference filters were used - green (544/6 nm), yellow (578/10 nm) and red (701/10 nm). The control module used was the original Leitz-ploemopcunit with mercury lamp equipped with I2/3 - cube (Exitationfilter 450/90nm; Suppression filter 520nm); TRITC - cube (Exitation filter 545/30nm; Suppression filter 610/75nm) and N2 - cube (simplified cube with Suppression filter LP 590nm only). The control module was used only to evaluate DiI-stained neurons and fluorescent beads. The list and characteristics of all used parts for each of the microscope modules are given in Tab.2.

To demonstrate the possibility of connecting multiple lasers to a single optical fiber, a branched optical fiber with eight separate inputs and one common output was chosen (Fig.2.B); the fiber core diameter was 25 µm. This fiber was not used in the practical part of the work due to its small diameter and, consequently, an excessively small illuminated field (although there may be applications where precisely a small illuminated area would be advantageous). However, it illustrates the concept: by merely switching different lasers on/off and changing emission filters, one can achieve rapid transitions between different excitation wavelengths without the need to manually reconnect the fiber to a new laser each time, as was done in all cases in this work. The relative complexity lies only in sourcing a branched optical fiber of suitable diameter, as thick branched single-mode fiber is relatively rare and expensive (unlike its non-branched counterpart).

### 2.7. Additional Optical Devices and Modifications for Evaluating the Ellis Concept

To assess quantitative fluorescence characteristics (fluorescence quantum yield) when comparing the Ellis concept with classical mercury lamp fluorescence microscopy, the ФМЭЛ-1A microspectrophotometric attachment was used. Its fundamental optical schematic is shown in Fig. 3.A. The ФМЭЛ-1А is a tube mounted in place of the microscope’s optical head (Fig. 3.B). Inside the tube, at a distance of 105.5 mm from the reference plane (the surface by which the ФМЭЛ-1А attaches to the microscope body or the filter cube block), there is an aperture—an adjustable pinhole (with diameters of 1 µm, 5 µm, 15 µm). Beyond the pinhole is a series of filter slots (or an empty slot—position 0). Past the filters, a collimating lens expands the light beam exiting the pinhole and directs it onto the ФЭУ-79 (photomultiplier tube - PMT). The ФЭУ-79, in turn, is connected to auxiliary electronics in the following sequence: a resistor bank (1-10 MΩ) → У5-9 усилитель → Щ4300 voltmeter. The ФЭУ-79 is powered by the УБПВ-1 high-voltage power supply. The ФМЭЛ-1A allows for measuring fluorescence intensity at a defined point (quantitative data), limited by the pinhole diameter, via voltmeter readings. To nullify differences in ФМЭЛ-1А readings when comparing the two systems (Ellis vs. the classical Leitz Labovert fluorescence microscope), the same Leitz filter cubes from the original filter cube block were used in both cases (Fig.3.В); the ФМЭЛ’s own filters were set to position 0. The ФМЭЛ-1А optical system also includes a side optical path enabling real-time observation simultaneously with fluorescence intensity measurement. A more detailed description of the device is provided in the official manual.

**Figure 3.**
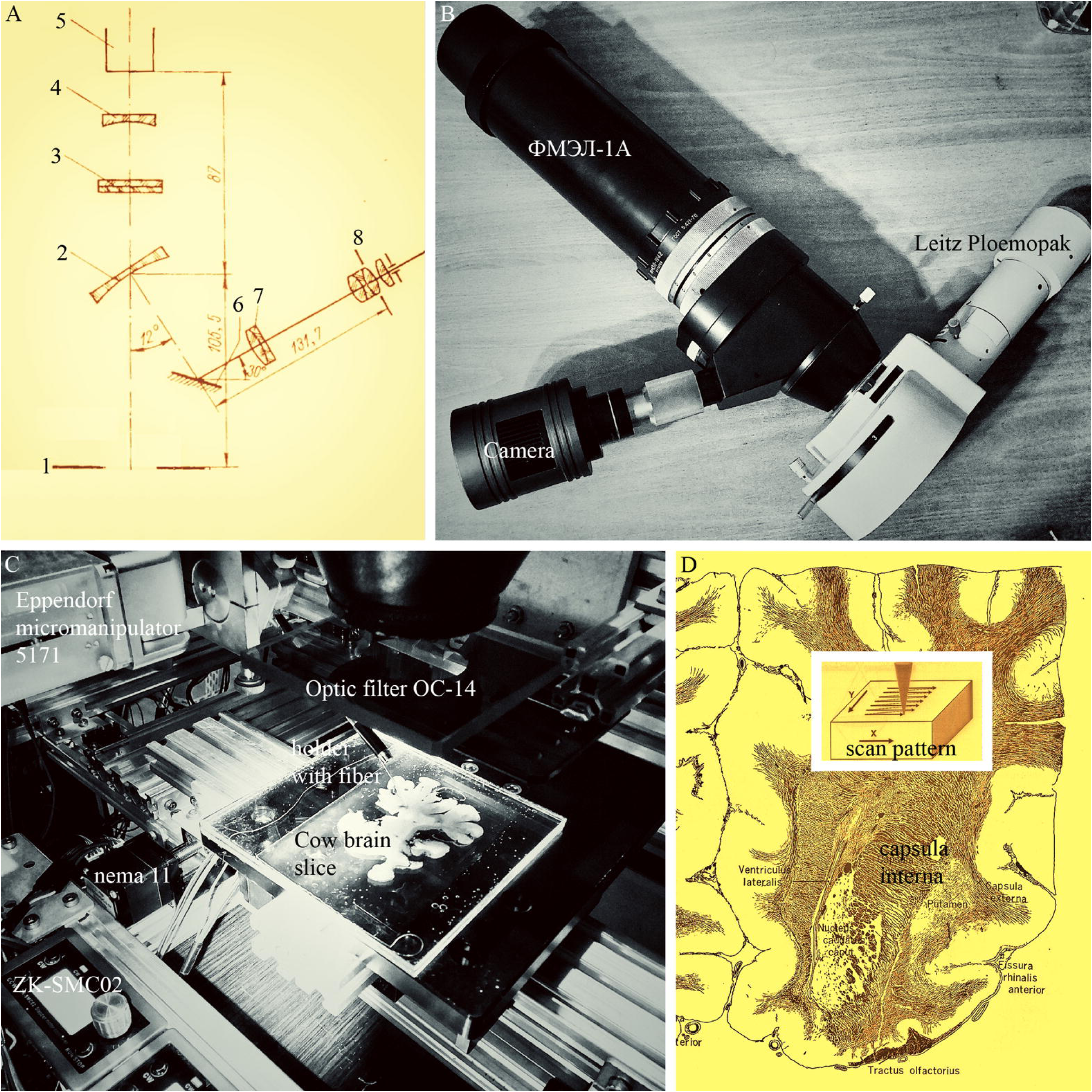
A - Optical diagram of the microphotometric attachment ФМЭЛ-1A (the picture taken from Микроскопы., 1969). The numbers in the figure indicate the following: 1 – the attachment base, used to mount it to the Ploemopak block instead of the microscope’s optical head (in the case of a Leitz Laborlux S); 2 – a mirror with a central pinhole; 3 – interference filters (in this work, these were replaced by the filter cubes of the Ploemopak block); 4 – a reverse lens, expanding the light beam passing through the pinhole; 5 – a photomultiplier tube (PMT); 6 – a mirror; 7 – lens of the side optical path; 8 – eyepiece. B – Photo of the ФМЭЛ-1А nozzle connected to the Leitz vacuum block and with a digital camera instead of an objective lens in the side optical path (the side path is the same microscope optical head, only integrated into the ФМЭЛ-1A). C – Photo of the Amscope 7x-45x stereomicroscope equipped with the Eppendorf 5171 electronic micromanipulator and a motorized stage based on linear translation modules with NEMA 11 motors, controlled via the ZK-SMC02 programmer. An ОС-14 glass optical filter is installed in front of the stereomicroscope objective. D – Scheme for scanning a large sample (calf brain slice) with an optical fiber secured on an electronic micromanipulator, implemented in practice on the stereomicroscope described above. The scheme is a combination of diagrams taken from Wright and Wright., 2002; Yoshikava, 1968). The calf brain slice analyzed in this work was taken at the rostral level of the capsula interna.

To demonstrate scanning capability using micromanipulators and confirm the universality of the Ellis concept, an AmScope 07-40 stereomicroscope was modified (Fig. 3.C). The design of this model does not allow for factory fluorescence modification; there is no filter cube block for it. However, the Ellis concept circumvents this problem; it is sufficient to add an emission filter in front of the stereomicroscope’s objective—in this case, an appropriately sized ОС-14 glass filter with a transmission range of 580-2700 nm (a filter for DiI). For scanning, the electronic micromanipulator Eppendorf 5171 was chosen, specifically its mechanical part with NEMA 17 stepper motors, each connected to a ZK-SMC02 programmer (Fig.3.C). The rest of the original 5171 manipulator electronics was unnecessary, as the ZK-SMC02 is more convenient. The programmer has 128 built-in programs for selecting the desired sample scanning mode. A steel holder with a single-mode optical fiber (1 mm core diameter) was mounted on the micromanipulator. Furthermore, a separate motorized stage based on NEMA 11 linear translation stages was created for the stereomicroscope (dual translation stages with 100 mm travel on the X-axis and dual translation stages with 200 mm travel on the Y-axis). Each pair of translation stages was connected to a separate ZK-SMC02 programmer. Aluminum structural frame parts were used as connecting nodes between the translation stages and to mount the micromanipulator. The motorized stage enables work with large, non-standard samples, such as calf brain slices (sample slide size: 100 mm × 100 mm), and the Eppendorf 5171 electronic micromanipulator has sufficient travel range along its axes (25 mm) for scanning a defined area on the slice (Fig.3.D).

### 2.8. 3D-vibrating microtome design description

At the heart of the basic concept of any vibratome model lies a blade that performs saw-tooth, reciprocating movements with a fixed amplitude along the x-axis. The sample advances towards the blade, which is fixed along the y-axis, and after the blade cuts through the sample, the sample returns back to its original position (along the y-axis). Then, the blade moves downward along the z-axis by a predetermined increment, and the cycle repeats. This work utilized a 3D vibratome (Klepukov, 2025) with the most advanced technical characteristics. The entire design of the vibratome can be divided into 4 modules (Fig.4.A).

**Figure 4.**
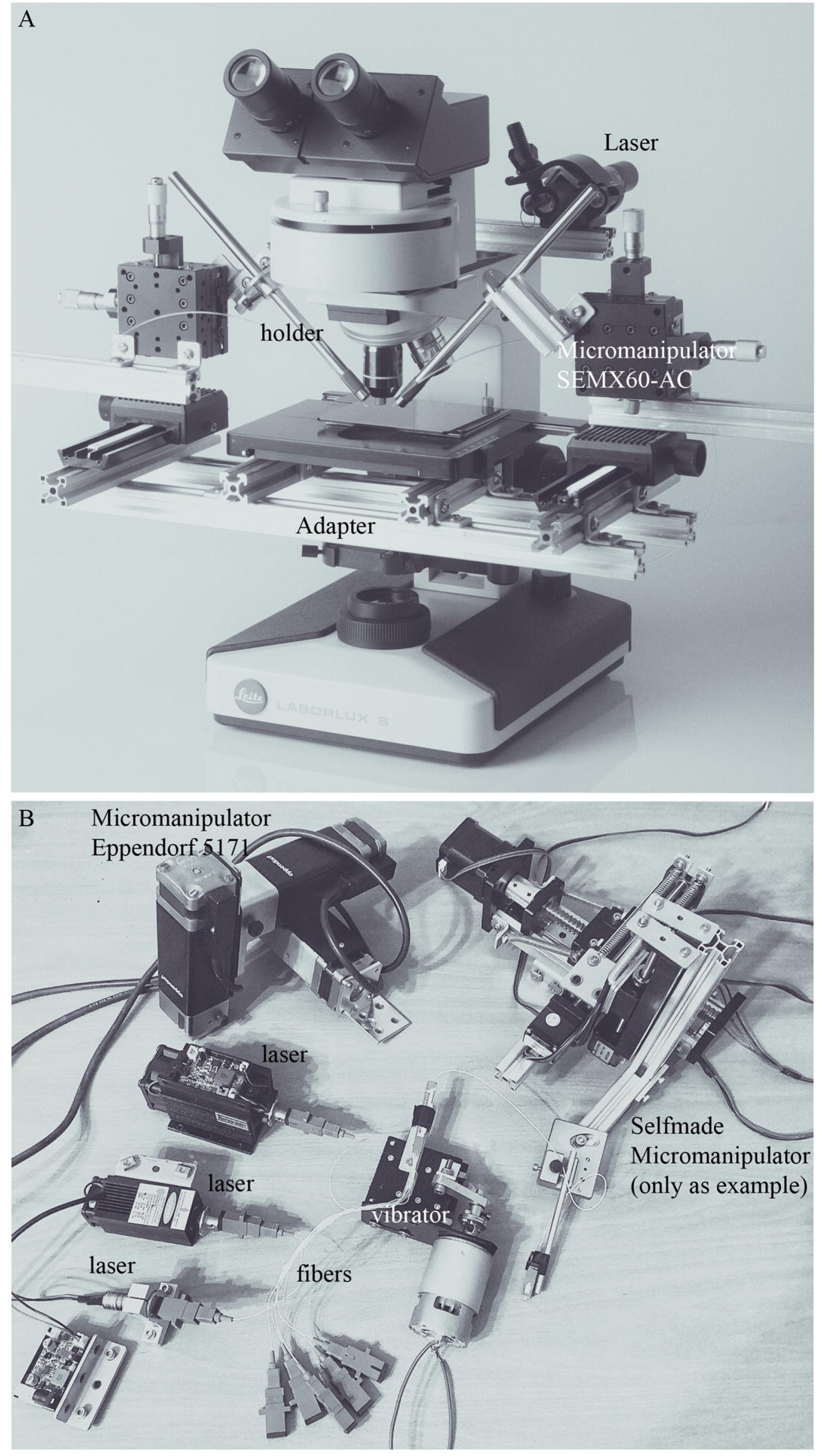
A - It is a front view of the 3D- vibrating microtome. Explanations of abbreviations for the aluminum profile and other parts are given in Table 3. B - electronic control module. It consists of three power supplies (24V, 2A, 50W) connected to three electronic programmers ZK-SMC02. Each of the programmers is connected to a stepper motor on the corresponding axis of the 3D-vibrating microtome (only y-axis in this case). Both power supplies and programmers are housed in an aluminium metal construction case. The programmer set is universal and allows controlling both the stepper motors of the 3D vibrational microtome and any electronic micromanipulator equipped with stepper motors (any type).

**Table 3.**
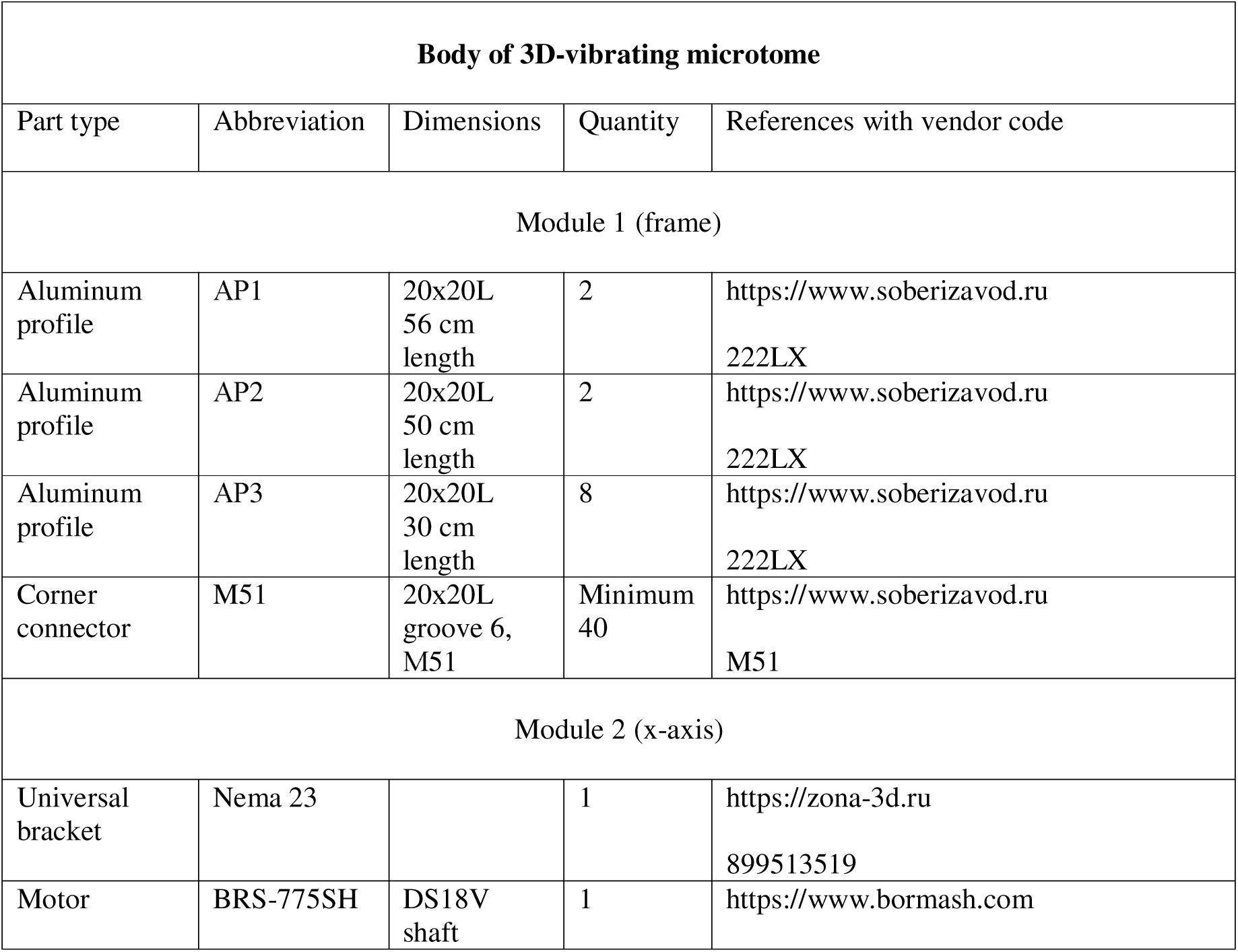
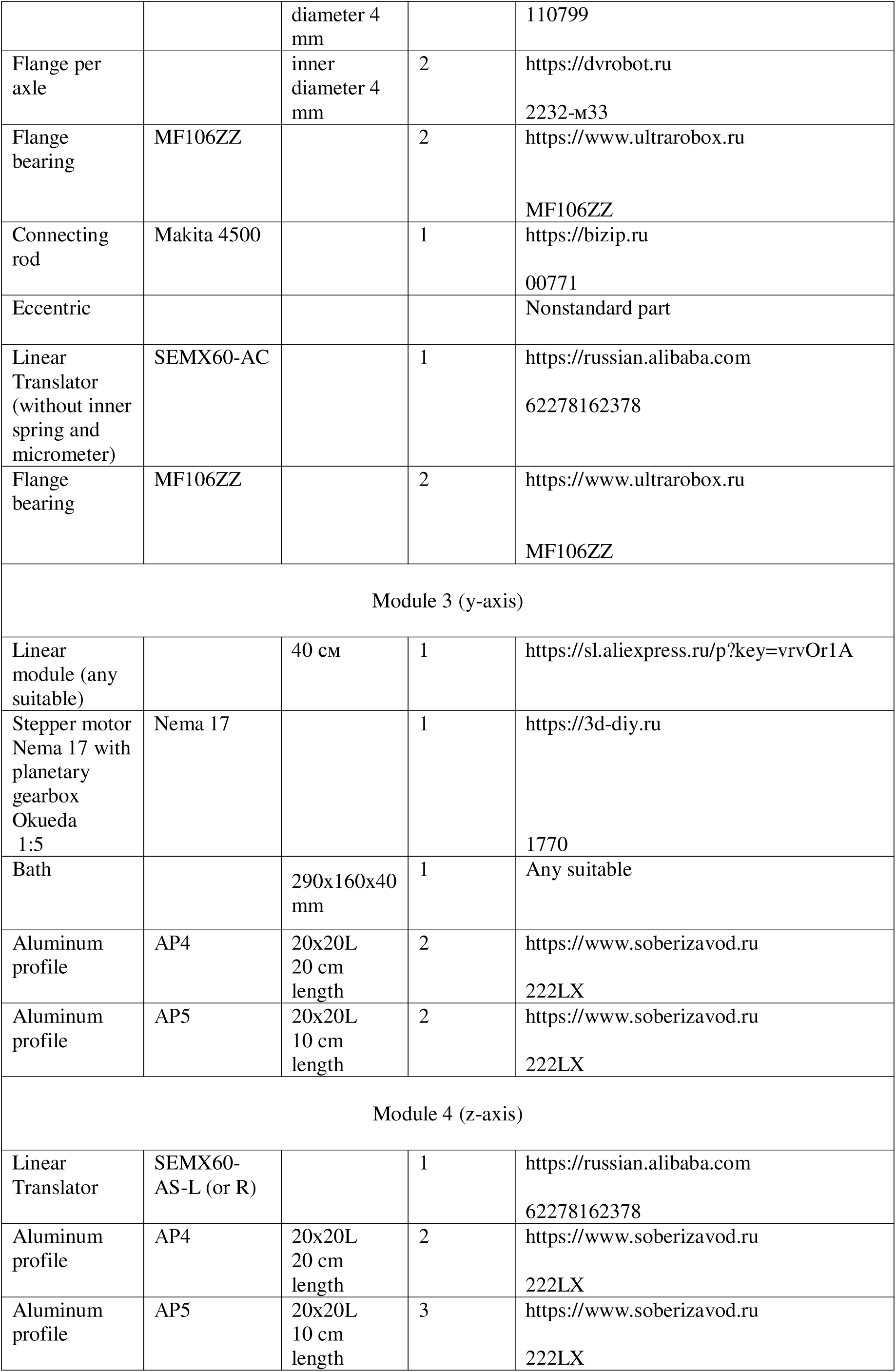
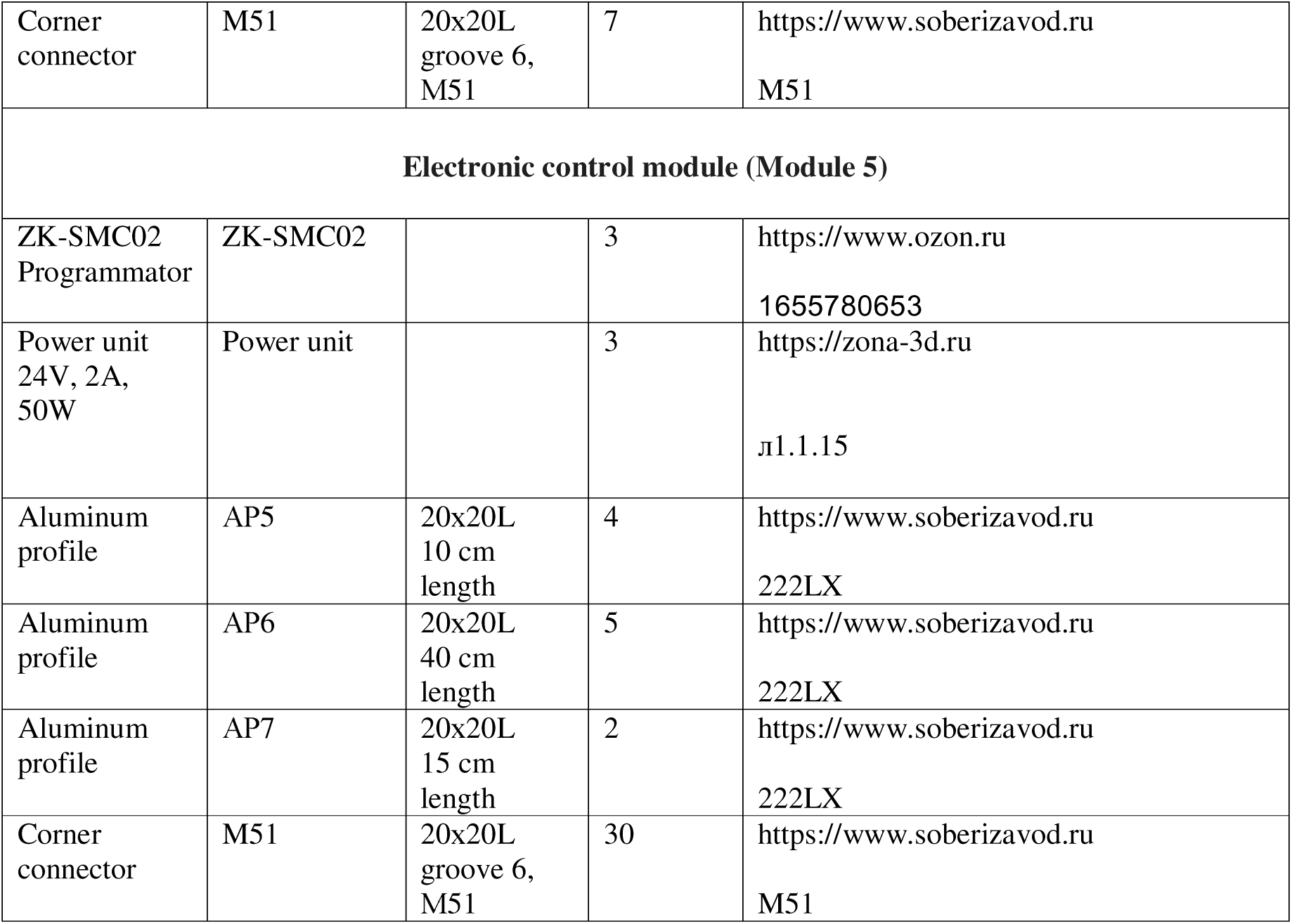
List of necessary parts of the corresponding abbreviations required to build a 3D-vibrating microtome. The list of parts is divided into modules that form the microtome.

**Table 4.** Image intensity values in pixels, calculated using ImageJ software. Values are given for three points for 10 μm beads.

| 10 μm beads<br>Light |  |  |  |  | 10 μm beads<br>Two lasers 532nm |  |  |  |  |
| --- | --- | --- | --- | --- | --- | --- | --- | --- | --- |
| Bead<br>intenci<br>ty | Galo<br>intensity |  |  | Backgrou<br>nd<br>noise | Bead<br>intenci<br>ty | Galo<br>intensity |  |  | Backgrou<br>nd<br>noise |
| 255<br>(same<br>for 10<br>beads) | 177 | 209 | 224 | 0.001<br>(same for<br>10 beas) | 255<br>(same<br>for 10<br>beads) | 231 | 220 | 186 | 0.001<br>(same for<br>10 beads) |
|  | 202 | 231 | 213 |  |  | 218 | 195 | 217 |  |
|  | 211 | 230 | 199 |  |  | 207 | 212 | 165 |  |
|  | 209 | 213 | 214 |  |  | 224 | 206 | 148 |  |
|  | 178 | 177 | 188 |  |  | 199 | 206 | 221 |  |
|  | 210 | 212 | 212 |  |  | 228 | 217 | 181 |  |
|  | 219 | 213 | 236 |  |  | 171 | 213 | 203 |  |
|  | 166 | 210 | 183 |  |  | 238 | 188 | 206 |  |
|  | 242 | 216 | 231 |  |  | 197 | 189 | 193 |  |
|  | 177 | 217 | 153 |  |  | 216 | 164 | 123 |  |
| Mean<br>255 | Mean 205.733 |  |  | Mean<br>0.001 | Mean<br>255 | Mean 199.4 |  |  | Mean<br>0.001 |

**Table 5.** Image intensity values in pixels, calculated using ImageJ software. Values are given for three points for 3 μm beads.

| 3 μm beads<br>Light |  |  |  |  | 3 μm beads<br>Two lasers 532nm |  |  |  |  |
| --- | --- | --- | --- | --- | --- | --- | --- | --- | --- |
| Bead<br>intenci<br>ty | Galo<br>intensity |  |  | Backgrou<br>nd<br>noise | Bead<br>intenci<br>ty | Galo<br>intensity |  |  | Backgrou<br>nd<br>noise |
| 255<br>(same<br>for 10<br>beads) | 193 | 139 | 154 | 0.001<br>(same for<br>10 beas) | 255<br>(same<br>for 10<br>beads) | 228 | 199 | 184 | 0.001<br>(same for<br>10 beads) |
|  | 237 | 216 | 200 |  |  | 195 | 236 | 143 |  |
|  | 233 | 185 | 180 |  |  | 224 | 237 | 211 |  |
|  | 183 | 228 | 235 |  |  | 187 | 242 | 173 |  |
|  | 223 | 158 | 201 |  |  | 180 | 240 | 171 |  |
|  | 211 | 203 | 220 |  |  | 196 | 220 | 154 |  |
|  | 239 | 244 | 197 |  |  | 174 | 215 | 204 |  |
|  | 231 | 216 | 200 |  |  | 198 | 178 | 235 |  |
|  | 217 | 214 | 196 |  |  | 234 | 210 | 194 |  |
|  | 176 | 217 | 211 |  |  | 149 | 218 | 165 |  |
| Mean<br>255 | Mean 205.233 |  |  | Mean<br>0.001 | Mean<br>255 | Mean 199.8 |  |  | Mean<br>0.001 |

To assemble the “First Module” - the load-bearing frame for the other three modules - parts from an aluminum construction set were used (Tab.3). The “Second Module” of any vibrational microtome is the mechanism that provides the reciprocating motion of the blade along the x-axis (a crank-and-connecting rod mechanism). In this mechanism, an eccentric is attached to the electric motor (BRS-775SH, DS18V, 4mm shaft diameter) via two flanges. A connecting rod (Makita 4500) is mounted onto the eccentric via bearings (MF106ZZ). The other end of the connecting rod is connected, via two bearings (MF106ZZ) mounted on an axle (a screw of suitable diameter), to a platform movable along the x-axis (an opto-mechanical platform SEMX60-AC with the micrometer removed and without the internal spring). The motor transmits motion to the platform via the connecting rod, and the eccentric ensures the platform is displaced by a specified amount (determined by the offset in mm between the axes of the eccentric). The electric motor is started by directly supplying it with direct current from a standard laboratory power supply. The motor and the SEMX-60 are attached to the load-bearing frame via a bracket and parts from the aluminum construction set.

“The third module” is a self-contained linear translator. Its core is a platform mounted on linear bearings, which is driven by the rotation of a ball screw (BS). The ball screw is connected via an adapter to a stepper motor, such as a Nema 17. A planetary gearbox (Okueda) is installed between the ball screw and the stepper motor, reducing the ball screw’s rotation speed by a specified ratio (1:5). This is a convenient feature because the lower the feed rate of the sample, the smaller the achievable section thickness. The stepper motor is controlled via a ZK-SMC02 programmer (Fig.3.B). A removable tissue sample bath is attached to the linear translator platform using an aluminum construction set (e.g., T-slot framing).

“The Fourth Module” is a mechanism that provides micrometric movement of the blade along the Z-axis. The vibratome implements a design in which an identical optomechanical platform, SEMX60-AR (in its original, full configuration and with a micrometer on the right), is attached at a 90-degree angle to the “second module” — the SEMX60-AC platform (without the internal spring). The blade itself is then attached to this SEMX60-AR platform via an adapter made from an aluminum construction set and a 40×40 connector with a double angle bracket. The design of the adapter allows for the blade angle to be adjusted over a wide range. For cutting small samples, a trapezoidal Dexter SK5 blade with a cutting edge length of 60 mm is suitable.

### 2.9. Making microparticle slides for fluorescence microscope calibration

For any new concept for a lighting system in microscopy, the most important parameter is how well this system can evaluate a point object, such as fluorescent beads with a diameter of ∼200 nm (Rigor Science; ∼ adsorption peak is 450-490 nm). To obtain a preparation with fluorescent beads (200 nm), 10 µl of the original stock solution (1%) was placed between two coverslips (each coverslip 0.17 mm thick).

Furthermore, to assess the signal-to-noise ratio, fluorescently stained particles with diameters of 3 µm and 10 µm (Jingsu Zhichuan Technology) were prepared. The original stock solution contains unstained particles, and to evaluate fluorescence, the particles need to be stained with a fluorescent marker. DiI was used for staining. One DiI crystal (10 µm in diameter) was mixed with 50 µl of the original particle stock solution (particle concentration in both cases: 2.5%) and 50 µl of technical acetone. After the DiI crystal was completely dissolved, 10 µl of the resulting mixture was mixed with 90 µl of distilled water. The final concentration of the DiI-stained beads was 0.125%. To obtain a preparation with fluorescent beads (3 and 10 µm), 10 µl of this solution (0.125%) was placed between two coverslips (each 0.17 mm thick). It is important to note that DiI is very poorly soluble in water; therefore, its own concentration in the final solution is negligible (all DiI settles on the beads) and does not affect the signal-to-noise ratio.

### 2.10. USAF 1951 test and determination of the illuminated area

To evaluate the resolution of the fiber-optic microscope, a USAF 1951 test slide with a minimum resolution of 2.2 µm (group 7, element 6) was used. The evaluation was performed at 20x magnification, illuminating the slide with two 532 nm lasers and using a Leitz E3 filter cube.

In addition to the resolution test, measurements were taken using a micrometric scale to assess the illumination area that could be achieved using micromanipulators. Furthermore, using the ФМЭЛ-1A instrument, fluorescence intensity measurements were taken at different points within the illuminated area on a calibration slide (included with the ФМЭЛ-1A). The calibration slide is a special luminescent plate made of uranium glass ЖС19. When setting up the microscope with this 1.5 mm thick calibration slide, the microscope is focused on a crosshair marked on the slide’s surface. For intensity measurements, the image of the crosshair is moved outside the pinhole aperture to measure the fluorescence intensity.

## 3. Results

### 3.1 Brief description of the brain structure under study and its connections

For convenience in assessing the quality of the fiber-optic fluorescence microscope, a well-studied subject, Habenula (Hb), was chosen; it is a paired epithalamic nucleus consisting of two parts, Medial Habenular nucleus (MHb) and Lateral Habenular nucleus (LHb); the latter nucleus consists of two parts, Lateral part (LHbL) and Medial pert (LMbM). In rodents, the MHb is connected by afferent connections (Herkenham and Nauta., 1977) with the Triangular Septal nucleus (TS); the LHb is connected with the Lateral Preoptic area (LPA); afferents come as part of the bundle of stria medullaris (sm). Both TS and LPA show stained neurons when lipophilic dyes are applied to the MHb or LHb, respectively, and these connections are non-overlapping (if application to the MHb stained neurons in the LPA, for example, this suggests diffusion of the marker from the MHb to the LHb; Klepukov and Apryshko., 2019). In addition, the MHb is connected by efferent connections with the interpeduncular nucleus (IP), the efferents go as part of the compact fasciculus fasciculus retroflexus (fr); the LHb also gives efferents to the Midbrain via fr (Herkenham and Nauta., 1979).

By making labels of different lipophilic fluorescent markers in Hb, one can observe a beautiful distribution pattern of marker-labelled fibers and neurons on vibrational slices of mouse brain. Since the markers fluoresce at different excitation wavelengths, it becomes possible to test the widest possible set of lasers. The only limitation is that it is inadvisable to use lenses with magnifications greater than 50x, as larger magnifications are uninformative in the case of macro-assessment of neuron and fiber distribution.

### 3.2. Scanning of a Large Sample Using a Fiber Optic. Demonstration of Peripheral Illumination by a Fiber Optic on DiI-Stained Structures at Different Magnifications (Novelty of the Method)

Unlike the plomeopak block, illumination via the fiber optic is delivered not from above (through the objective) but from the side. This characteristic defines the entire subsequent workflow. Firstly, the illumination area achieved by the fiber optic at low magnifications (up to 20x) is smaller than when using a plomeopak block. Secondly, the fiber optic is initially mounted on a micromanipulator, which means that to achieve maximum brightness in a small illumination area, the fiber optic must be positioned correctly.

Micromanipulators are the key component. They allow for very precise orientation (down to ∼160 nm in the case of the Eppendorf 5171) of the fiber optic at any point on the slice, even if it is a non-standard size, such as a large calf brain slice (Fig. 5A). The initial assessment of such large objects is meaningful only at low magnifications under a stereomicroscope (overview view). The travel range of the micromanipulator axes allows for scanning across the entire surface of the slice (Fig.5B,С), especially if the manipulator is electronic and programmable (automatic scanning). Scanning enables finding all stained areas on the slice and illuminating them at low magnification (Fig. 5D-I). The stereomicroscope model is irrelevant here, as using an aluminum construction set allows the micromanipulators and emission filters to be mounted on any stereomicroscope. Subsequently, knowing the location of a specific marker-stained region on the large slice, it can be examined more precisely at higher magnifications directly under a fluorescence microscope.

**Figure 5.**
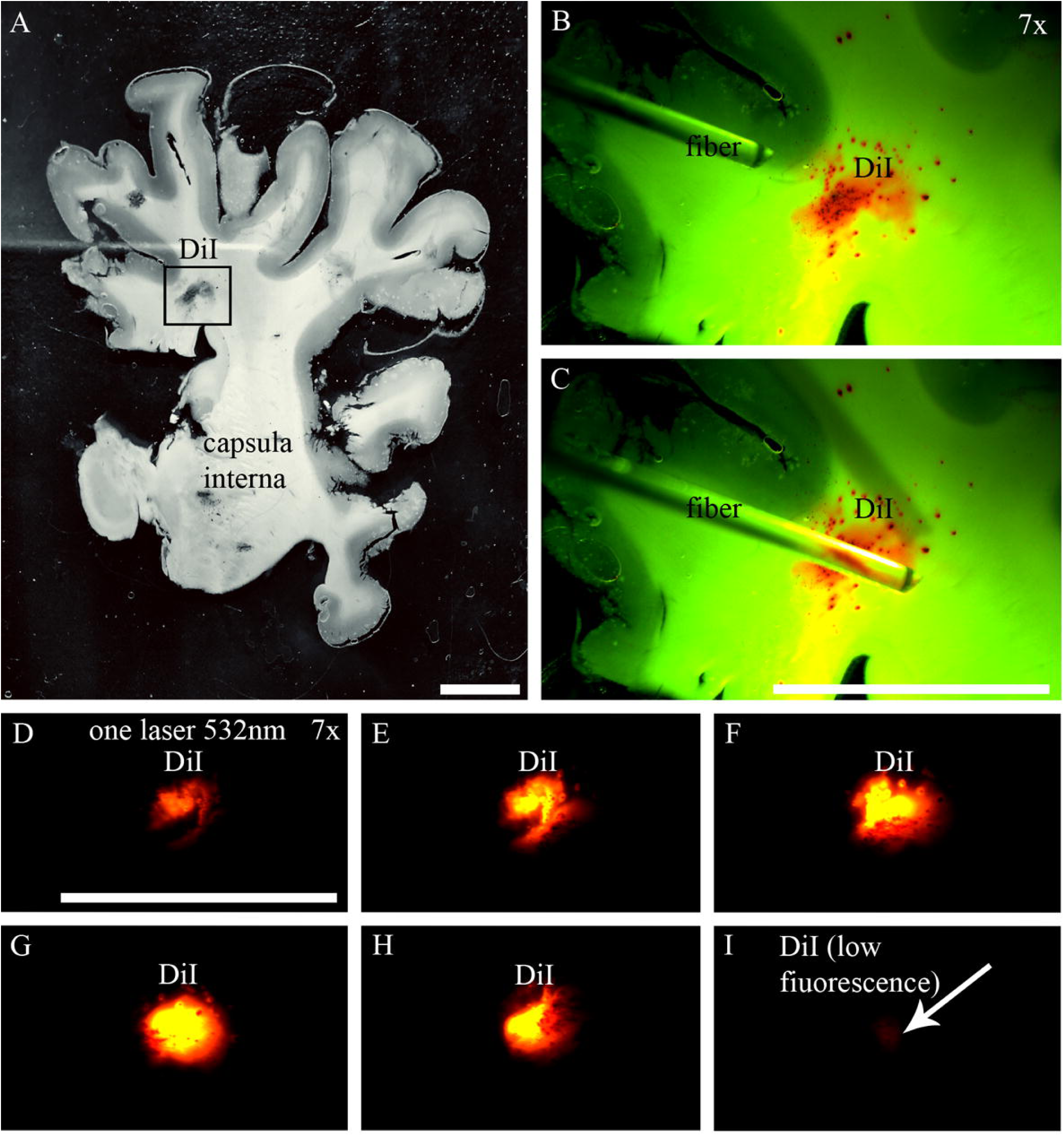
A - Overview image of a vibratome slice of a calf brain at the level of the rostral capsula interna (slice thickness 150 µm) with DiI crystals applied to the surface in different locations, soaked in technical acetone (for better solubility, black frame). B, C - Image of an optical fiber under the stereomicroscope objective, positioned using the Eppendorf 5171 micromanipulator over different areas of the slice (outlined with squares on the overview image) and illuminated with transmitted light. D-I - Scanning of DiI crystals with the optical fiber under fluorescent light (using a single 532 nm laser for illumination) along the slice surface with an ОС-14 filter. The optical fiber cannot illuminate the entire slice at once due to its small diameter, and therefore scanning the slice is the only way to detect structures labeled with the marker at very low magnifications, in order to subsequently examine them at higher magnification in a targeted manner. Scanning serves here as a search mechanism. First photo was taken by Ipad10. All other photos were taken with a Sony IMX178 sensor. Bar - 1 сm.

Micromanipulators also play a decisive role in evaluating stained objects at high magnifications (up to 50x). Typically, to achieve maximum illumination brightness of an object, the fiber-illuminated area needs to be positioned at the center of the frame. If the illuminated area is, for example, at the edge of the field of view, it needs to be carefully moved to the center. During such precise positioning, an interesting effect can be observed—as the illuminated area constantly changes, different parts of the object are illuminated differently. Ultimately, if desired, one can achieve either maximum illumination of the sample or illumination of only a small part of it, depending on the position of the illuminated area, by adjusting only the micromanipulators. With a plomeopak block, this is possible only by narrowing the field diaphragm, which sharply limits the field of view.

This capability – illuminating small portions of an object using a micromanipulator – was clearly demonstrated using a DiI-labeled Habenula structure on a frontal mouse brain slice (case 1, unilateral marker injection into the central part of LHb) at various magnifications (Fig. 6). At all magnifications, the optical fiber (in this case, just one) was initially positioned on the left border of the field of view and was moved to the right using the micromanipulator, gradually illuminating more and more structures within the slice. Upon reaching the central position, the fluorescence brightness became maximal and then decreased again as the fiber passed further to the right. The effect of varying fluorescence intensity depending on the position of the illuminated area was evident at 4x magnification (Fig.6.A-H), 10x (Fig.6.I-P), and 20x magnification (Fig.6.Q-X; at 50x it is more convenient to evaluate individual neurons). At 4x, for example, a single fiber can only illuminate a small labeled area of the MHb, while the main remaining part of the LHb remains non-fluorescent (as it is outside the illuminated field). Such selective illumination is impossible when using a planepack block, which can only change the overall field of view (via the field diaphragm).

**Figure 6.**
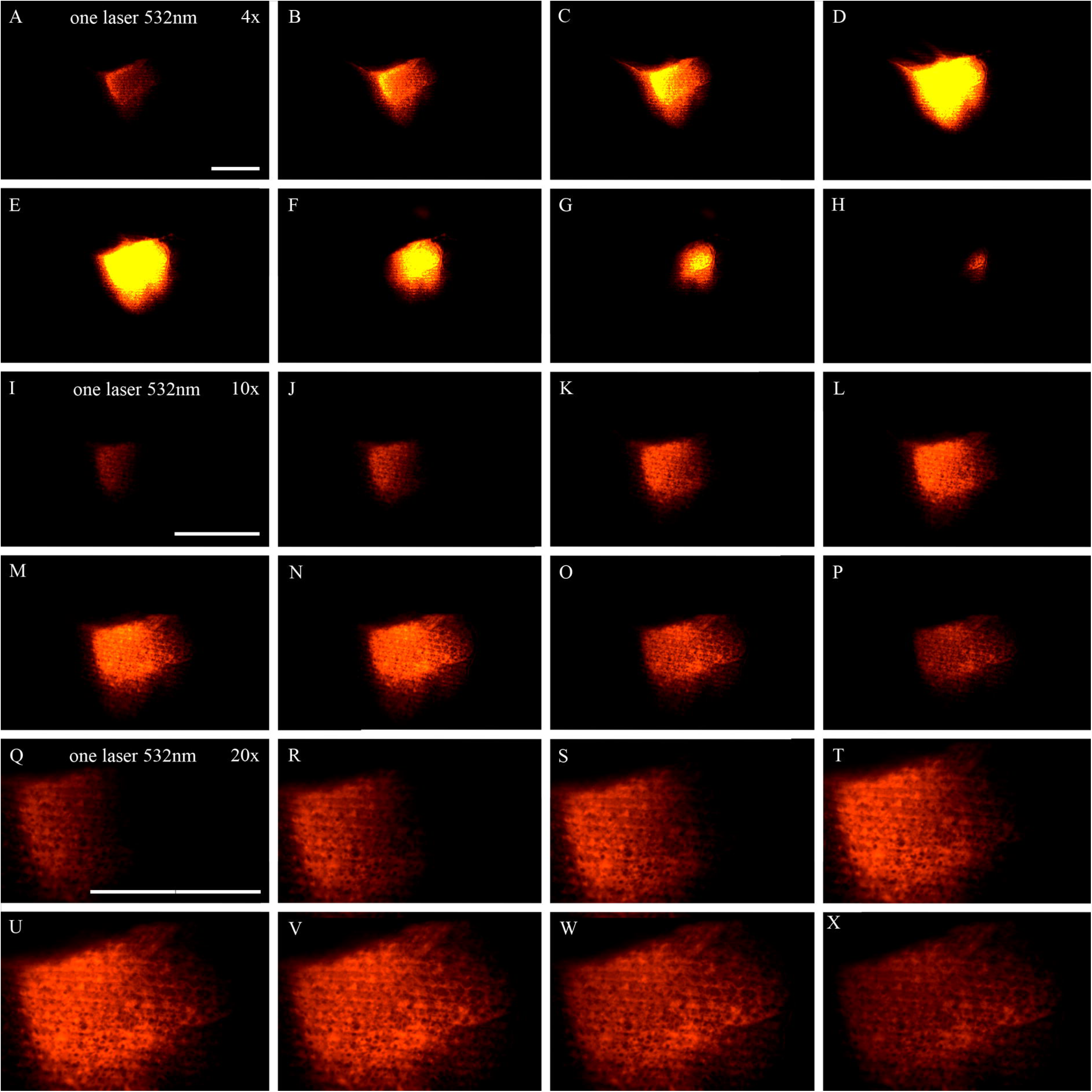
A – H - Sequential scanning with a single optical fiber (1mm diameter; 532 nm wavelength, TRITC filter) of a 100 µm thick mouse vibratome slice at the level of the Habenula (Hb), 4x magnification, DiI staining (crystals are localized in the central part of the lateral nucleus). At the periphery of the row (A, H) only a minor part of the slice is illuminated, with maximum brightness achieved in the center of the row (D,E) when the illuminated area from the optical fiber is concentrated on the crystals. Sequential movement of the optical fiber along the slice illuminates specific areas of the slice as chosen by the researcher. I-P - Sequential scanning with a single optical fiber of the same mouse vibratome slice, but at 10x magnification. Q-X - Sequential scanning with a single optical fiber of the same mouse vibratome slice at 20x magnification. It is important to emphasize that the effect of illuminating different areas on the slice is achievable at different magnifications. In the first photo of the series (Q) approximately a quarter of the total surface area of the stained region is illuminated at peak intensity (U,V). All photos were taken with a Sony IMX178 sensor. Bar – 100μm.

The example described above clearly demonstrates the direct influence of micromanipulators on which structures within the slice will be revealed. If desired, using two micromanipulators, one can illuminate different areas of the object within the field of view, changing the relative positions of the optical fibers and adjusting the illumination level of specific areas at will, while still keeping the entire object under illumination. This is unattainable with a planepack block, where light statically falls only from above through the objective, and the light flux can only be limited by the field diaphragm or ND filters. This constitutes the novelty of the technology. This capability was demonstrated on the same mouse Habenula structure (Fig. 7A-D), with the difference that in this case, instead of a 100 µm thick vibratome slice, the illuminated target was the surface of a cut made in a whole, unsliceed brain (case 2) where DiI had been injected (the brain was used as an example; the same would be visible on a slice).

**Figure 7.**
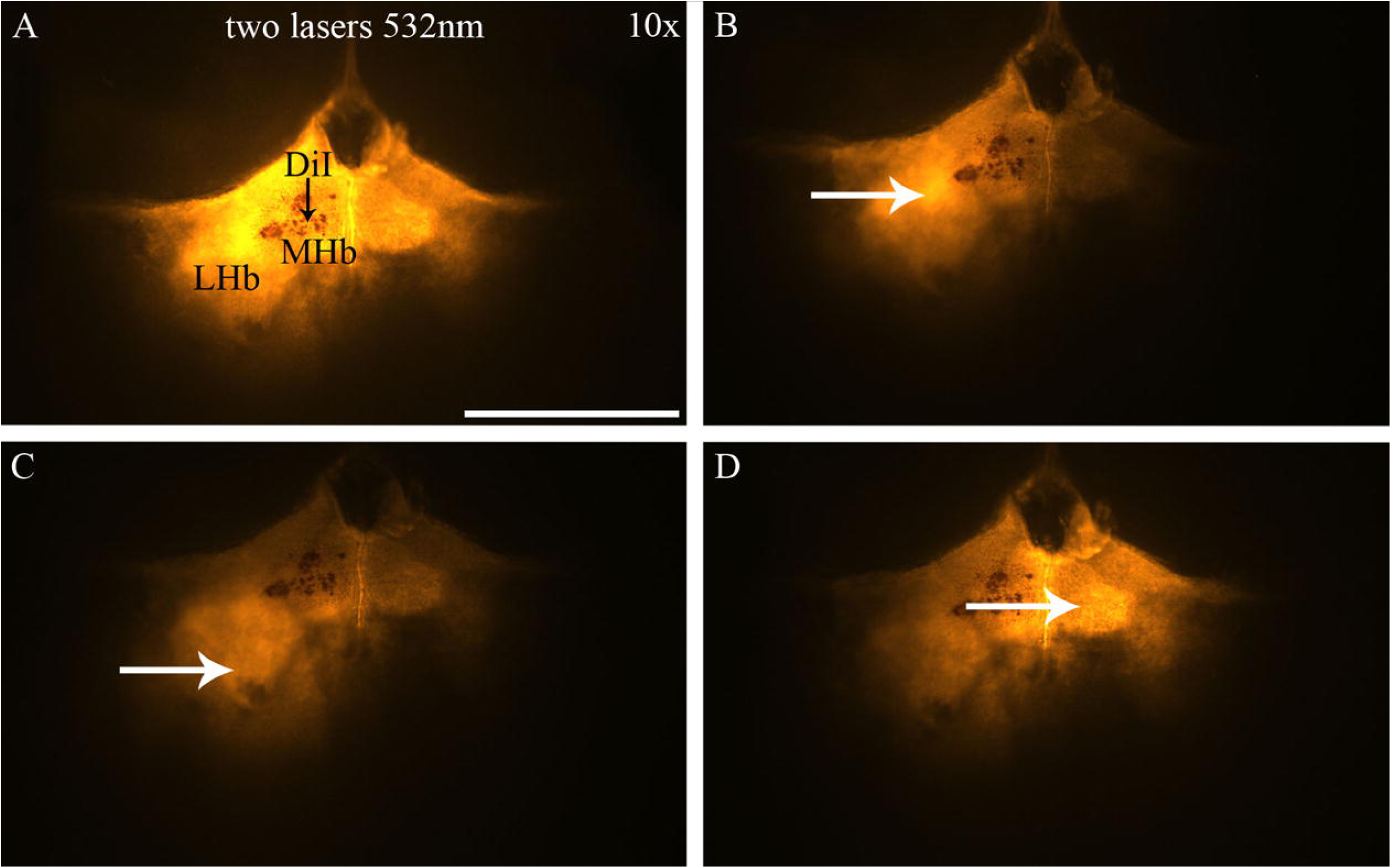
Illumination of the entire surface of a mouse brain with two optical fibers (dimeter 1 mm), featuring a large DiI label at the border between MHb and LHb on the left side of the brain (where fluorescence is brightest). A - The optical fibers are oriented to maximize the illuminated area of the slice. B - The optical fibers are shifted so that the maximum illumination falls on the central-left area of the LHb (white arrow), while the right side of the brain is barely illuminated. C - The optical fibers are shifted so that only a small area in the lower-left quadrant is illuminated (white arrow). D - The optical fibers are adjusted for maximum illumination of the central brain region between the MHb nuclei of adjacent hemispheres (white arrow); meanwhile, the LHb of the left hemisphere fluoresces weakly, despite the presence of individual large DiI crystals there. All photos were taken with a Sony IMX178 sensor. Bar - 100μm.

### 3.3 Comparison of Fluorescence Quantum Yield under Mercury Lamp vs. Optical Fiber Illumination at Different Magnifications. Evaluation of a Brain Sample with DiI Injections

The fluorescence quantum yield is defined as the ratio of the number of emitted photons to absorbed photons. The quantum yield indicates the efficiency of the fluorescence process. The better the object is illuminated, the higher the fluorescence quantum yield (efficiency), and the more details can be observed. A quantitative measure of quantum efficiency can be provided by electrometer readings (in millivolts) obtained from the ФМЭЛ-1A microspectrophotometric attachment. The object used for evaluating the mercury lamp system (Fig.8А,С,Е) vs. the Ellis concept system (Fig.8.В.D.F) was a mouse brain slice at the level of the Hb (the same slice as in the results above). Illumination was provided by two lasers (532 nm) and two corresponding optical fibers positioned opposite each other (the object was placed at the interslice of the light from both fibers). The assessment of fluorescence quantum yield was performed using 4x, 10x, 20x, and 50x objectives with a TRITC filter cube. The pinhole diameter on the ФМЭЛ-1A was 1 µm, the amplifier (Усилитель У5-9) was set to three, the resistance before the amplifier was 1 MΩ. The background current value was 0 mV. The results of the quantum yield measurements in millivolts are as follows (accounting for background current):

Mercury lamp light vs. optical fiber (two fibers) illumination:
For the 4x Leitz EF objective: 10.2 mV vs. 11.2 mV;
For the 10x LMPlan objective: 10 mV vs. 10 mV;
For the 20x LMPlan objective: 10 mV vs. 7 mV;
For the 50x LMPlan objective: 7 mV vs. 3.5 mV.

**Figure 8.**
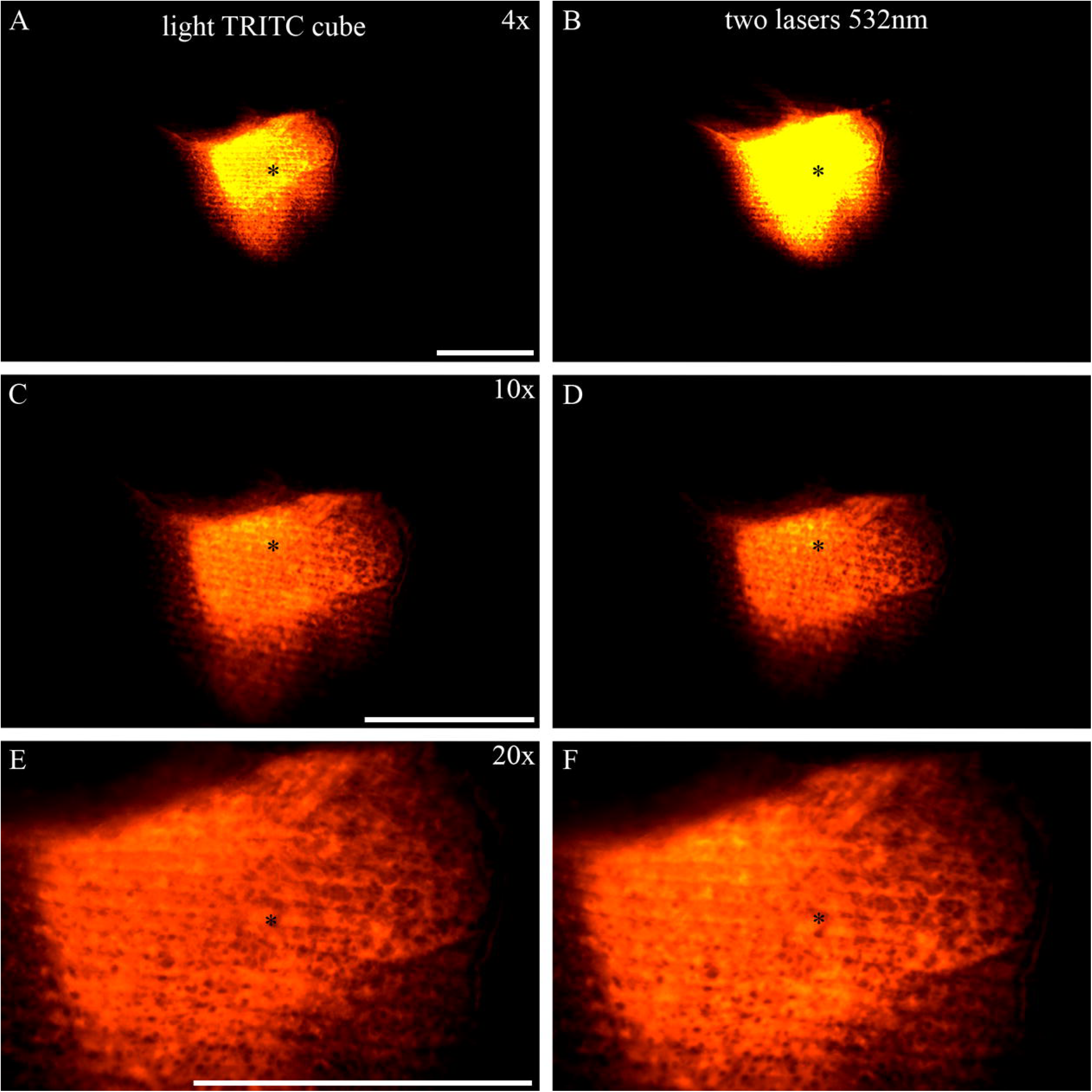
A, C, E - Illumination with mercury lamp light through the vacuum block (TRITC filter) of a DiI-stained area on the same mouse brain vibratome slice as in Fig. 6. Objectives 4x (A), 10x (C), and 20x (E) were used. At different magnifications, quantum yield of fluorescence (in mV) was measured using the ФМЭЛ-1A attachment (pinhole diameter 1 µm, little white star). For the Leitz EF 4x objective, this value was 10.2 mV; for the LMPlan 10x objective - 10 mV; for the LMPlan 20x objective - 10 mV; for the LMPlan 50x objective - 7 mV (for 50x no photo was taken). The lower the objective magnification, the more photons from the illuminated area reach the ФМЭЛ-1A, resulting in a higher quantum yield value. B, D, F - Illumination with two optical fibers (532 nm, TRITC filter) of the surface of the same DiI-stained area at the same three different magnifications. For the Leitz EF 4x objective (B), this value was 11.2 mV; for the LMPlan 10x objective (D) - 10 mV; for the LMPlan 20x objective (F) - 10 mV; for the LMPlan 50x objective - 3.5 mV (for 50x no photo was taken). The numbers show that at 4x magnification under laser illumination (B), the quantum yield is even slightly higher than under mercury lamp illumination (A), which is also evident from the presence of some overexposure in the laser case (images were taken at equal camera exposure levels). At 10x, the quantum yield values are equal (C, D; equal camera exposure level). At 20x, the quantum yield value is lower under laser illumination (F) compared to mercury lamp illumination (E), but by adjusting the digital camera exposure values, images of equal brightness can be achieved (photos E and F were taken at different exposure values, adjusted manually). All photos were taken with a Sony IMX178 sensor. Bar - 100μm.

As can be seen from the figures, at low magnifications (4x), the quantum yield is slightly higher with optical fiber illumination, which is also noticeable during visual comparison of the slice with mercury lamp illumination (Fig. 8.A.B). Even some excessive illumination is noticeable with fiber optic lighting, despite the fact that imaging was performed at the same exposure value in both cases. At 10x magnification, the quantum yield values equalize (Fig.8.C.D). At 20x (Fig.8.E.F) and 50x magnifications, the quantum yield is higher with mercury lamp illumination (for 50x, only measurements were taken, no photo). However, by adjusting the camera exposure value, this difference can be negated (the 20x image under laser light (Fig. 8.F) was taken at a slightly higher camera exposure to demonstrate comparable image quality even with different quantum yields).

### 3.4. Evaluation of brain sample with DiI injections (without Leitz-cube)

Another case (3) with DiI tag was also evaluated to assess the way to improve the quality of the obtained images. Here a different camera was used (Allied Vision Marlin F145B2), slice thickness of 50 µm (vs. 100 in the case above) and two green lasers (532nm). An only Zeiss emission filter (578/10) was used instead of a TRITC-filter cube. In this case the marker was localized in the MHb, but part of it also entered the hippocampus (Hi; Fig.9.A). The stained fasciculus retroflexus (fr) fibers were present in significant numbers, and viewed separately (Fig.9.B). The ventro-medial wall of the hippocampus revealed dense clusters of stained neuronal bodies, which were very clearly visualized due to parasitic DiI application (Fig.9.C).

**Figure 9.**
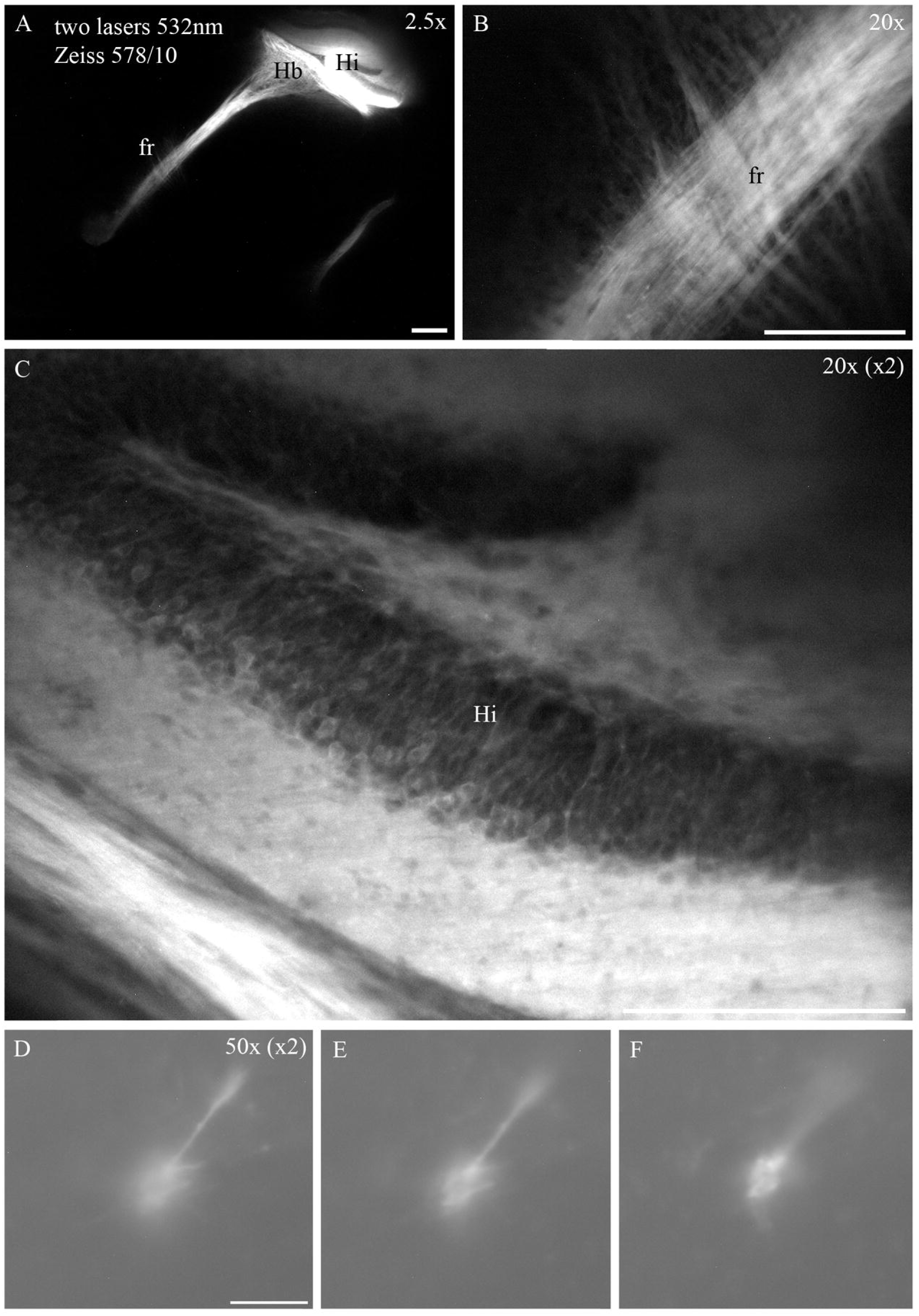
Improving image quality in a fiber optic lighting system is possible through a combination of factors. Firstly, two fibers can be used in a lateral position (or more) - the brighter the lightning of the object, the more detail is revealed. Secondly, the slice thickness can be reduced, this reduces the overall level of background fluorescence outside the focal plane of the lens. Finally, the use of sensors with a larger pixel size gives a deeper and more contrasty image. All three of these conditions were met when evaluating case 6, illuminated using two green lasers and fiber optics, which helped to maximize the level of detail. A - analysis of the distribution of DiI-labeled structures on sagittal vibrational slices of case-6 mouse brain (sag); the brain sample was cut into 50 μm-thick slices (on average this is 2 times lower than the standard thickness values for vibratory slices). A - DiI application was made through a sagittal section of the brain surface in MHb (Leitz EF 4x lens); the marker is localized in the dorso-caudal region of the nucleus (A, marked by arrows) and bordering the Hippocampus (Hi). Bar - 100μm. B - the large, MHb-associated axonal tract fasciculus retroflexus (fr) is perfectly visible, individual fibers can clearly be distinguished in fr (LM Plan 20x lens). Bar - 100μm. C - image magnified by 2 times compared to the original image of the labeled DiI bodies of neurons in Hi. The presence of stained neuronal bodies is solely due to the parasitic diffusion of the marker from the application site (MHb) into Hi. Nevertheless, this section of the brain sliceis very demonstrative, as it shows perfectly stained neuronal bodies despite their dense concentration (LM Plan 20x lens). Bar - 100μm. D,E,F – images magnified by 2 times compared to the original images of a single neuron near Hi at three different levels along the z-axis (step - 10 µm, photos were taken at high magnification using LM Plan 50x lens). These images were obtained to demonstrate the operation of the fiber optic system at high magnifications. Bar - 50μm. All photos were taken with a Sony ICX204 sensor.

Both fr and neurons in the hippocampus were imaged at 20x magnification. A single neuron (near the hippocampus) was specifically imaged at high magnification (50x) at three different levels along the z-axis (Fig 9.D,E,F). This is not a standard magnification for morphological work with DiI, but in this case it was necessary to evaluate the performance of the illumination system with this lens.

### 3.5. Evaluation of brain samples with B3-PPC injections (without Leitz-cube)

A total of three sagittal brain samples were used with B3-PPC injected into the MHb in a jelly-like semi-liquid form (case 4-6). The marker was localized within the MHb and partially in the LHb in all three cases studied. All mappings were exceptionally accurate and the entire MHb was represented in all three brain samples. The most representative case is shown in Fig.10A-C (case 5). Two 488-nm lasers and a Carl Zeiss emission narrowband interference filter (544/6nm) were used to visualise the B3-PPC labelled structures. The slices were evaluated at lens magnification of 2.5x, 10x. It should also be said that all these slices were also examined using a proprietary Ploemopak block with a mercury lamp and Leitz I2/3 filter cube (Exitation filter BP450-490nm; Suppression filter LP 515nm), but the resulting image was too dark even at high camera exposure values (data are not shown).

The application site was clearly visible and had a brighter colour compared to the surrounding MHb tissue (Fig.10.A). Fiber bundles in fr were clearly visible (Fig.10.B). No labelled neuronal bodies or fiber bundles were detected in the TS. Part of the marker also entered the LHb, which revealed numerous fiber bundles in the Midbrain (Fig.10.C). Dense bundles of labelled fibers in the sm were detected at the entrance to the dorsal LPA, but no visible neuronal bodies were also detected.

**Figure 10.**
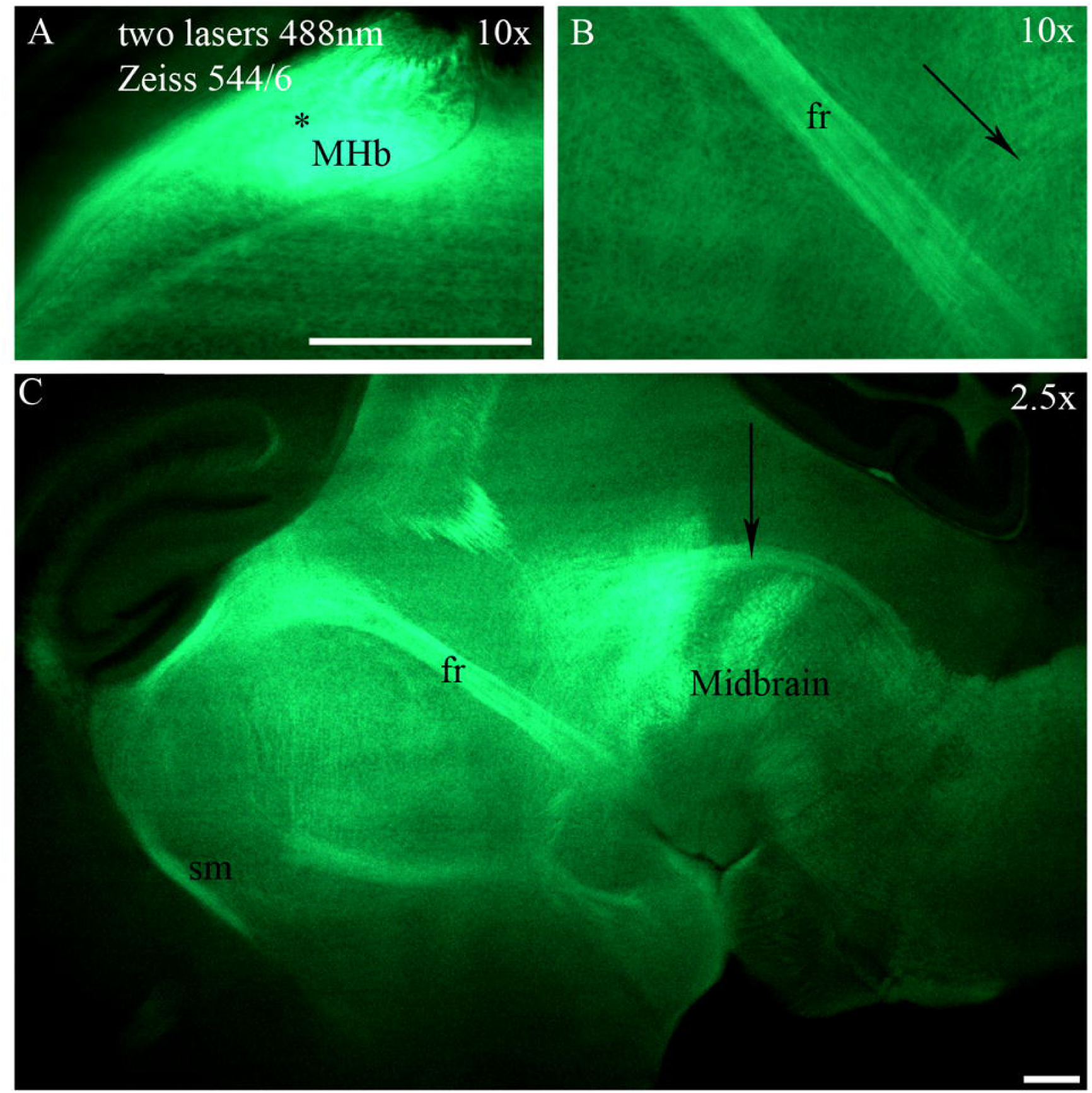
Distribution of B3-PPC-labeled structures in mouse brain P11 on sagittal slices. Illumination by two lasers at 488nm with fiber optics in the top position. Carl Zeiss emission interference filter (544/6nm) was used. A - location of B3-PPC staining in Medial Habenula (MHb), the marker is concentrated predominantly in the caudal part of the nucleus (marked with an asterisk). Magnification is 10x. B - image of a compact, perfectly stained with marker fasciculus retroflexus (fr) tract consisting of axon bundles emanating from the MHb. The caudal part of the tract shows scattered fibers (marked with an arrow) arranged perpendicular to the main fr fiber array. Magnification is 10x. C – Double magnified overview image of the brain slice at the level of the LHb. Stria medullaris (sm), fr and numerous strands of arc-shaped marker-labelled fibers in the Midbrain (marked with an arrow) are perfectly visible. No marker-stained neuron bodies were identified in the specimen. Magnification is 2.5x. All photos were taken with a Sony IMX178 sensor. Bar - 100μm.

Since no stained neuronal bodies were found in all three brain samples in the TS (and that is why three brain samples were taken - to set reliable statistics), but axonal tracts associated with the MHb were perfectly stained - all this allows us to potentially classify B3-PPC as a selective membrane tracer that stains only axons, but not neuronal bodies with their dendrites.

### 3.6. Evaluation of brain sample with DiD injections (without Leitz-cube)

One P11 mouse brain sample was used for DiD marker injection (case 7). The marker was injected in crystal form, unilaterally into the LHb (the sagittal surface of the cut while searching for the right level passed just lateral to the midline of the brain, and the MHb did not enter this sample). Two 650 nm lasers and a Carl Zeiss emission narrowband interference filter (701/10nm) were used to visualise the marker-labelled structures. The slices were evaluated at lens magnification of 2.5x, 10x.

The application site was clearly visible and almost completely filled with marker (Fig.11.A). The LHb had a brightly stained fr, from which numerous scattered fibers descended caudally perpendicular to the main trunk (Fig.11.B). The rostral afferent tract sm was also perfectly identified, with a maximum concentration of fibers in the caudal part, which was clearly visible by the intensity of the luminescence (Fig.11.C). Since the application was made in the LHb (the MHb was completely cut off), the TS neurons were naturally not detected, proving once again that the TS is innervated only by the MHb.

**Figure 11.**
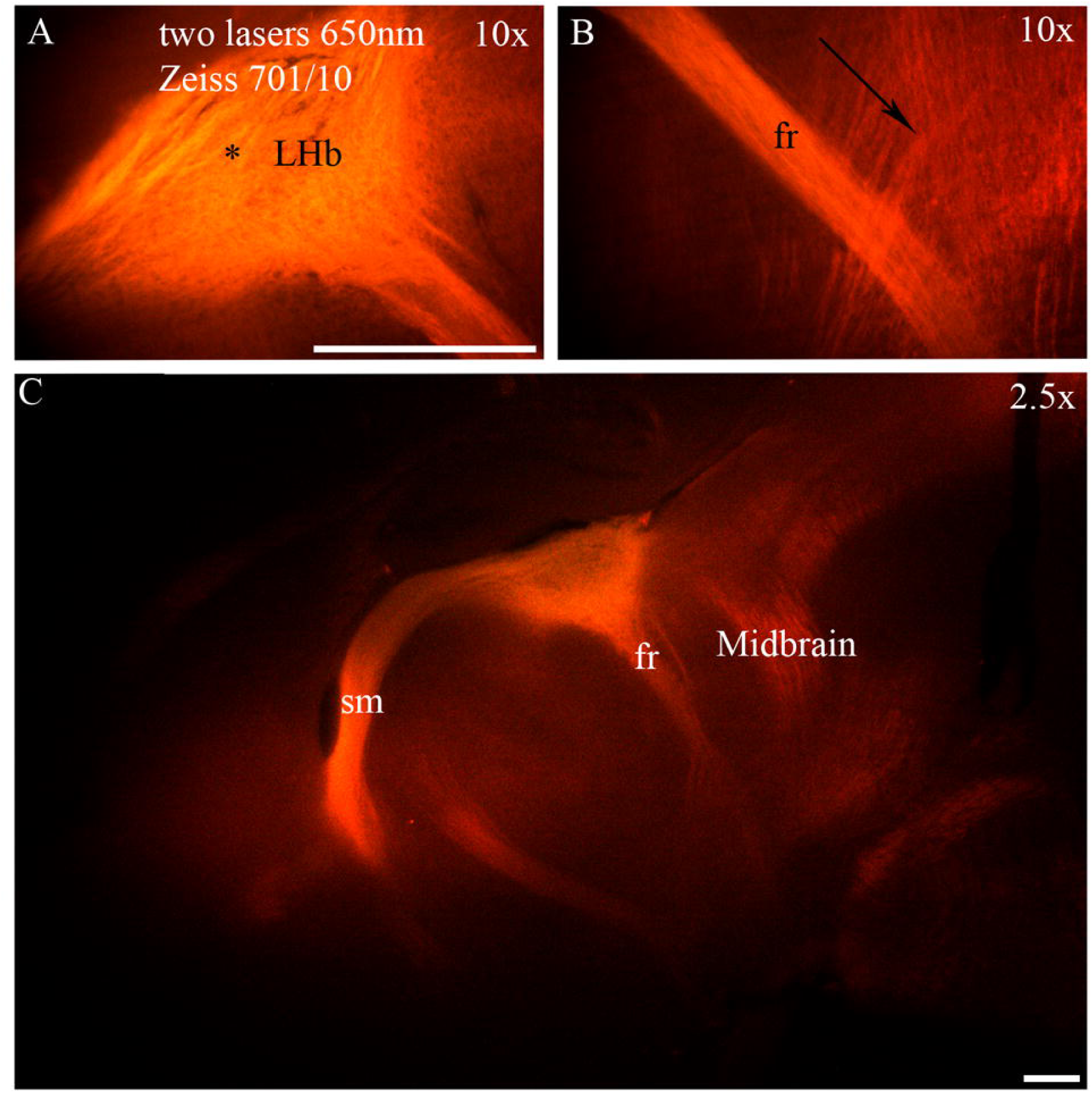
Distribution of DiD-labeled structures in mouse brain P11 on sagittal slices. Illumination by two overhead lasers at 650nm with fiber optics in the top position. A Carl Zeiss emission interference filter (701/10nm) was used. A - in this brain sample, the Medial habenula (MHb) was completely sheared off during level finding, so the application was made to the Lateral Habenula (LHb) on the sagittal surface. The application itself was accurate (marked with an asterisk) and the marker occupied most of the nucleus. Magnification is 10x B - image of a compact, excellent marker-stained fasciculus retroflexus (fr) tract consisting of axon bundles emerging from the MHb. The caudal part of the tract shows scattered fibers (marked with an arrow) emerging from fr and arranged perpendicular to the main fr fiber array. Magnification is 10x C - doubled overview image of the brain slice at the level of LHb. Excellent views of fr and especially the stria medullaris (sm), with maximum fiber density in the caudal part of the tract. Magnification is 2.5x. All photos were taken with a Sony IMX178 sensor. Bar – 100μm.

It should be noted that the shooting in the far red region of the visible spectrum took place at the maximum values of camera shutter speed, the picture in the eyepieces was almost indistinguishable to the naked eye (which is natural, since the human eye poorly perceives light near the infrared spectrum).

### 3.7. Assessment of Illuminated Area Size and Quantitative Measurements of Quantum Yield Across the Field of View

To evaluate the size of the illuminated area (in micrometers), a micrometer ruler and a luminescent plate made of ЖС19 glass (included in the ФМЭЛ-1A kit) were used. The plate was illuminated using a single optical fiber (Fig.12.A; core diameter 1mm), two optical fibers (Fig.12.B-H; core diameter 1 mm) with 488nm lasers, and using a ploemopak block. The assessment of the illuminated area was performed at 4x magnification using a Leitz EF objective and an E3 filter cube; measurements were taken against the micrometer ruler.

1. When using a single laser (Fig.12A), the length of the illuminated area along the x-axis was 230 µm (a), and along the y-axis 140 µm (b). The shape of the illuminated area is an ellipse. The area of the illuminated region is calculated using the formula:

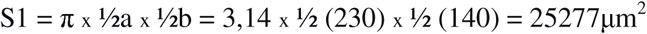

**Figure 12.**
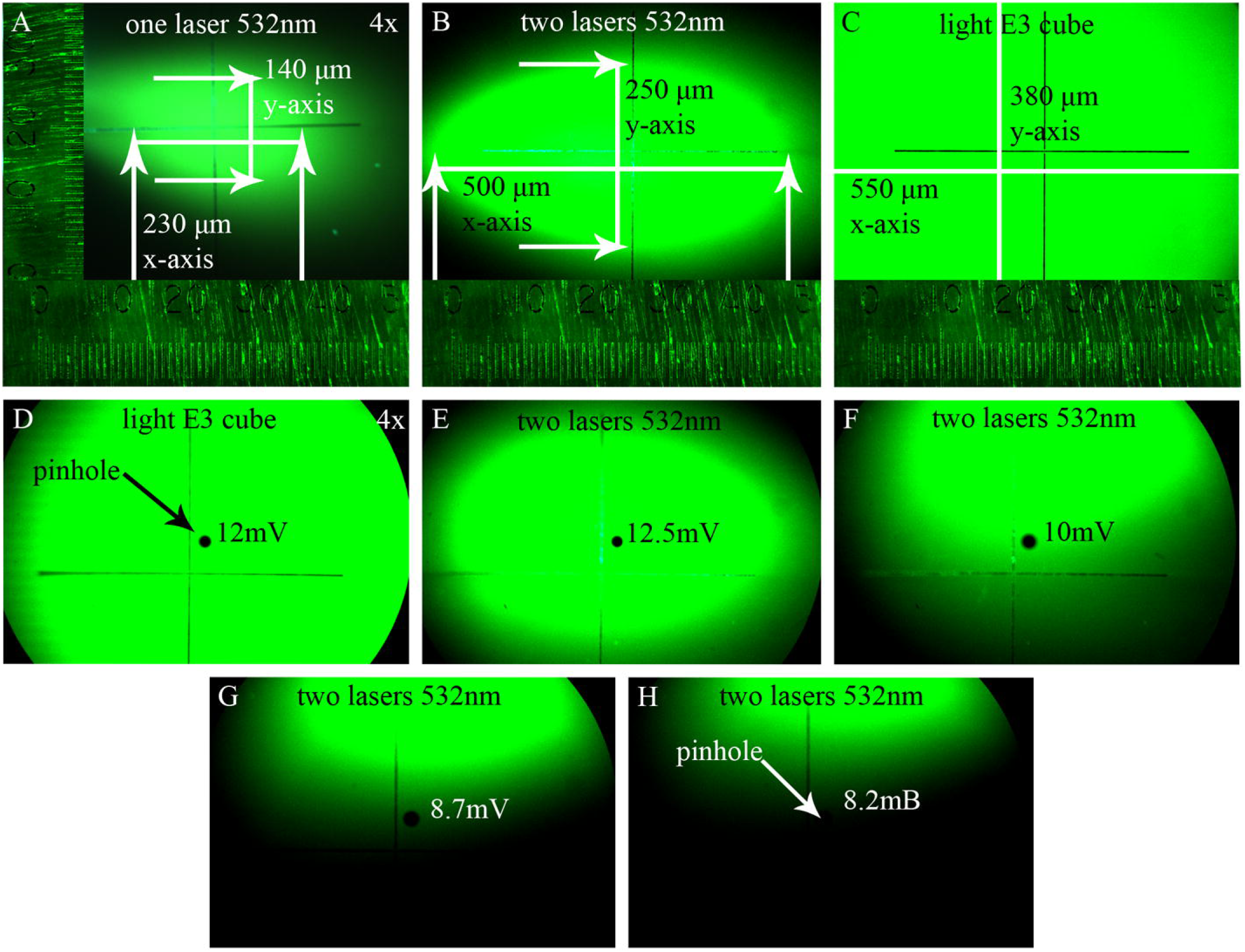
Evaluation of the illuminated area size was performed using a ЖС19 luminescent plate with a 4x Leitz EF objective and an E3 filter cube. Illumination was provided by one (A) or two (B) 488 nm lasers and by a mercury lamp (C). Measurements were taken using a micrometer scale (insets in the photos). A - When using one laser, the length of the illuminated area along the x-axis was 230 µm, and along the y-axis 140 µm. The shape of the illuminated area is an ellipse. The area of the illuminated region was 25277μm^2^ (see results 3.7). B - When using two lasers, the length of the illuminated area along the x-axis was 500 µm, and along the y-axis 250 µm. The shape of the illuminated area is an ellipse. The area of the illuminated region was 98125μm^2^ C - A control measurement of the illumination area with the mercury lamp showed that the entire area within the field of view was uniformly illuminated (see figure). The length of the illuminated area along the x-axis was 550 µm, and along the y-axis 380 µm. The shape of the illuminated area is a rectangle. The area of the illuminated region was 209000μm^2^ In addition to assessing the illuminated area on the ЖС19 luminescent plate, measurements of fluorescence quantum yield were conducted using the ФМЭЛ-1A microspectrophotometric attachment (D-H), with a pinhole diameter of 5 µm (black circle in the center of the frame). D - When illuminating the plate with a mercury lamp, the quantum yield value in mV was 12 mV. E - For two lasers at the center of the illuminated area - 12.5 mV. F - At the edge of the visible illuminated area - 10 mV. G - Beyond the illuminated area, approximately ∼70 µm from the boundary of the bright field - 8.7 mV. H - Beyond the illuminated area, approximately ∼140 µm from the boundary of the bright field - 8.2 mV. Based on the ФМЭЛ-1А readings, it can be concluded that the object is illuminated by the optical fiber not only within the visible boundaries of the field of view but also beyond this boundary, albeit to a lesser extent

Where π = 3.14, а is the length along the x-axis, b is the length along the y-axis.

2. When using two lasers with overlapping illuminated areas, the length of the illuminated region along the x-axis was 500 µm, and along the y-axis 250 µm. The shape of the illuminated area is an ellipse. The area of the illuminated region is calculated as:

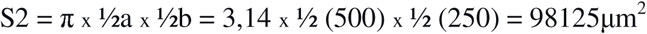
3. A control measurement of the illuminated area using a mercury lamp showed that the entire area within the field of view was uniformly illuminated (Fig.12.C). The length of the illuminated area along the x-axis was 550 µm (a), and along the y-axis 380 µm (b). The shape of the illuminated area is a rectangle. The area of the illuminated region is calculated using the formula (in µm²)

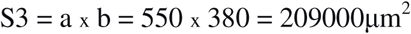

As can be seen from the obtained figures, when illuminating with a single laser, the illuminated area S1 constitutes approximately (25277/209000)x100=12.064% of the area illuminated by the mercury lamp (S3); when illuminating with two lasers (S2), it is (98125/209000)x100 = 46.94%.

In addition to assessing the illuminated area on the ЖС19 luminescent plate, further measurements of fluorescence quantum yield were conducted using the ФМЭЛ-1A microspectrophotometric attachment (Fig.12.D-H). The measurement parameters were: amplifier gain (Усилитель У5-9) set to three, resistance 1 MΩ; background current value 7.5 mV. The assessment was performed using an E3 filter cube. Digital camera imaging was conducted at the same exposure level with a 4x Leitz EF objective.

Measurements were taken both under mercury lamp illumination and under illumination with two 488 nm lasers (to achieve maximum field of view brightness; values for a single laser were therefore not measured). In the case of laser illumination, the pinhole (5 µm diameter) was positioned at the center of the illuminated area, at its edge, and beyond its edge (micromanipulators allow free control over the movement of the illuminated area within the field of view when using laser/fiber optic illumination). The quantum yield values in mV were:

- For the mercury lamp: 12 mV (Fig.12.D; value constant across the entire field of view).
- For the laser at the center of the illuminated area: 12.5 mV (Fig.12.E)
- For the laser at the edge of the illuminated area: 10 mV (Fig.12.F).
- For the laser approximately 70 µm beyond the boundary of the bright field: 8.7 mV (Fig.12.G)
- For the laser approximately 140 µm beyond the boundary of the bright field: 8.2 mV (Fig.12.H).

As evidenced by the obtained data, the quantum yield values at the center and edge of the laser-illuminated area do not differ significantly from (and are even slightly higher than) those obtained with the mercury lamp. What is more interesting is that the ФМЭЛ-1A detected weak fluorescence even at a distance of 140 µm from the edge of the visually illuminated area. This indicates that, at the specific exposure level used for the digital camera, the actual boundaries of the illumination field were wider than those perceptible to the human eye and covered the entire field of view.

### 3.8. Image Quality Assessment (Signal-to-Noise Ratio) Using ImageJ

Fluorescently pre-stained DiI beads with diameters of 3 and 10 µm were imaged at magnifications of 4x, 10x, 20x, and 50x using the epi-illumination block (TRITC filter; Fig.13.A,C,I) of two green 532 nm lasers (laser voltage 4.8V; Fig.13.B,D,F). The obtained images show visualization of 100% of the beads for each diameter, indicating that the entire field of view at these magnifications is illuminated. This is particularly significant in the context of laser illumination, as even when illuminated by two lasers, the illuminated area on slide ЖС19 at 4x magnification constitutes only ∼51% of the area illuminated by the epi-illumination block (Fig.12.B). Nevertheless, measurements using FMEL-1А (see results. 3.7) and photos of the beads clearly demonstrate that the fiber optic illumination covers more than 51% of the illuminated area (these 51% only represent the peak illumination intensity), since all beads at the periphery of the field of view are also visualized.

**Figure 13.**
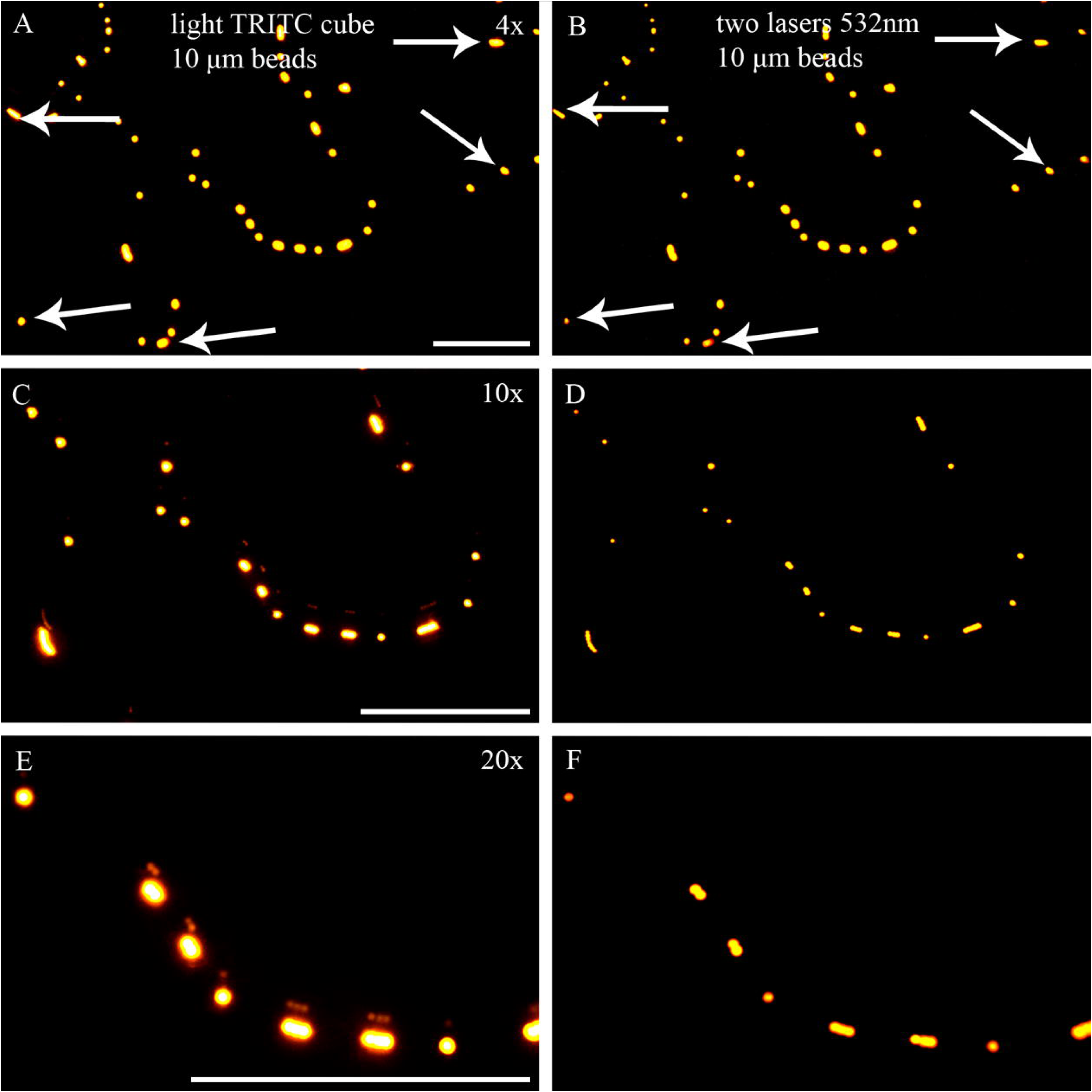
Comparison of the distribution pattern of pre-stained DiI fluorescent beads with a diameter of 10 µm at different magnifications (4x, 10x, 20x) under illumination with a mercury lamp light (A, C, E) and with two 532 nm lasers via optical fiber (B, D, F). At all magnifications, the number of detected beads was identical when using the two different systems, indicating that even at 4x (B) the entire field of view is illuminated by the laser, although at the periphery of the image it is noticeable that the fluorescence intensity of individual particles is lower (arrow) than in the center of the frame and compared to the same particles illuminated by mercury lamp light (white arrow). At higher magnifications, the difference in fluorescence intensity at the periphery of the field of view is almost imperceptible. All photos were taken with a Sony IMX178 sensor. Bar – 100μm.

To quantitatively characterize different illumination systems (mercury lamp light vs. laser light via optical fiber), images of pre-stained DiI beads with diameters of 3 µm and 10 µm at 50x magnification were analyzed (Fig. 14.A-D; for 200 nm diameter beads, the signal-to-noise ratio was not calculated due to their excessively small size). The ImageJ software was used to measure the intensity (in pixels) within the particle diameter, the intensity in the immediate vicinity of the particle (halo zone), and the background noise intensity (measured within one particle diameter from the particle’s size) for each image. Based on these three parameters, the signal difference-to-noise ratio (SDNR) and the contrast-to-noise ratio (CNR) were calculated for individual beads present in the captured images, using the corresponding equations (1), (2), and (3). The equations and the evaluation concept via ImageJ were taken from Sami et al., (2023):

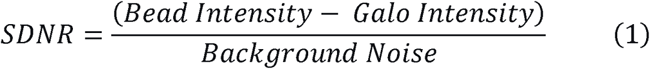

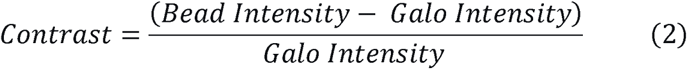

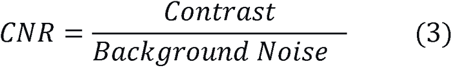

**Figure 14.**
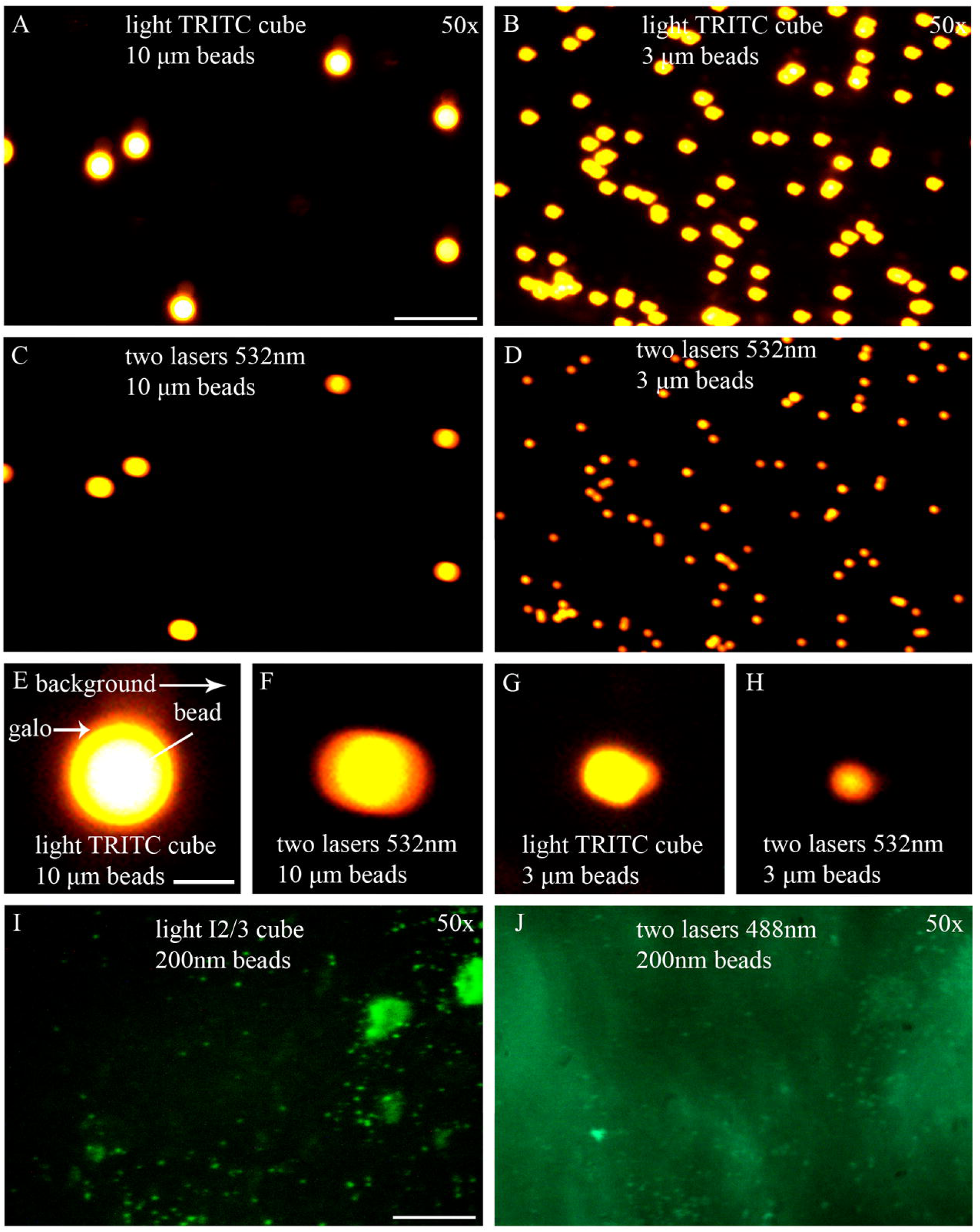
Comparison of pre-stained DiI beads with a diameter of 3 µm (A, B) and 10 µm (C, D), imaged at 50x magnification with different illumination systems (mercury lamp light vs. two 532 nm lasers via optical fiber (1 mm diameter)). The number of detected particles was identical under illumination with both systems, and at this high magnification it is convenient to measure fluorescence intensity in pixels (using ImageJ software). Bar – 50μm The measurement was taken at three points in three regions: in the center of the bead, in the halo, and at a distance of one bead diameter from the halo (white lines, E). Measuring intensity in pixels allows for calculating the signal-difference-to-noise ratio (SDNR) and contrast-to-noise ratio (CNR) for individual beads. Differences in halo size under mercury lamp and laser illumination (E-H) were noted: the other form of halo diameter created by lasers compared to illumination of the same bead by the mercury lamp. Bar - 10μm In addition, separately, without SDNR and CNR analysis (due to their small size), beads with a diameter of 200 nm were imaged (mercury lamp light (I) vs. blue 488 nm laser(J)). The fact that these beads were detected speaks to the resolving power of the fiber-optic illumination system, enabling it to capture particles of this size. Moreover, the use of lasers in this specific case is even more preferable, as these particular particles fluoresce best at a wavelength closer to 488 nm, and therefore the image obtained with laser illumination is brighter than with the fluorescent filter cube. Bar – 50μm

To determine the SDNR and CNR values for the entire image, the average SDNR and CNR values for ten beads within that image were calculated (Tab.4,5). The assessment was conducted using three points for each bead: three points within the bead itself, three points in the halo, and three points for the background (Fig.14.E). The following figures were obtained:

**10 µm beads, mercury lamp illumination:**

1. SDNR value: (255 - 205.733) / 0.001 = 49267
2. Contrast value: (255 - 205.733) / 205.733 = 0.239
3. CNR value: 0.239 / 0.001 = 239

**10 µm beads, illumination from two lasers via optical fiber:**

1. SDNR value: (255 - 199.4) / 0.001 = 55600
2. Contrast value: (255 - 199.4) / 199.4 = 0.279
3. CNR value: 0.279 / 0.001 = 279

**3 µm beads, mercury lamp illumination:**

1. SDNR value: (255 - 205.233) / 0.001 = 49 767
2. Contrast value: (255 - 205.233) / 205.233 = 0.242
3. CNR value: 0.242 / 0.001 =242

**3 µm beads, illumination from two lasers via optical fiber:**

1. SDNR value: (255 - 199.8) / 0.001 = 55200
2. Contrast value: (255 - 199.8) / 199.8 = 0.276
3. CNR value: 0.276 / 0.001 = 276

A higher signal-to-noise ratio (SDNR) indicates a cleaner signal with less interference. A higher contrast-to-noise ratio (CNR) indicates a sharper image. As evident from the figures, in both of these metrics (SDNR and CNR), the laser light via optical fiber slightly outperforms the mercury lamp light, with the laser values being somewhat higher.

Differences in halo size under mercury lamp and laser illumination (Fig.14.E-H) were noted: the other form of halo diameter created by lasers compared to illumination of the same bead by the mercury lamp. This interesting anomaly is likely related to the epi-illumination properties of the objective and its aperture diameter, as well as the fact that light falls on the beads at different angles in the different illumination systems. The halo is essentially the object’s shadow, and it will differ just as the size of a shadow differs when illuminated by the midday sun versus at sunset.

In addition to the 3 µm and 10 µm beads, 200 nm beads were also imaged using the epi-illumination block (I2/3 filter; Fig.14.I) and two 488 nm lasers (Fig.14.J). These beads possess intrinsic fluorescence (DiI staining is not required and is impossible as it overlaps with the spectrum) and are more sensitive to the 488 nm wavelength. For this reason, the number of beads visible under laser illumination is significantly higher than under illumination with the epi-illumination block equipped with a broadband filter (450-490 nm). However, in the context of this work, the number of detected beads is not the primary focus (for which purpose the DiI-stained beads are more convenient here), but rather the very fact of their detection, as this fact demonstrates the resolution capability of the microscope with laser illumination via optical fiber.

### 3.9. USAF 1951 Test Results

For the quantitative assessment of the achieved optical resolution, a USAF 1951 test target was used. Figure 15.A. shows an overview image of the USAF 1951 test target, with a detailed view (Fig. 15B,C) of element 6 in group 7, indicating an optical resolution of 2.2 µm. Element 6 of group 7 is the smallest element available on the specific USAF test target in the author’s possession. Furthermore, an image of the same target area was captured under conditions without fiber optic vibration (Fig. 15D). This image is significantly noisier (pixelated), serving as an excellent demonstration of the phenomenon of laser coherence, which fiber vibration is intended to eliminate. Figure 15.E.F shows a pixel intensities of vertical and horizontal lines of Group 7 Element 6 (made in ImageJ program).

**Figure 15.**
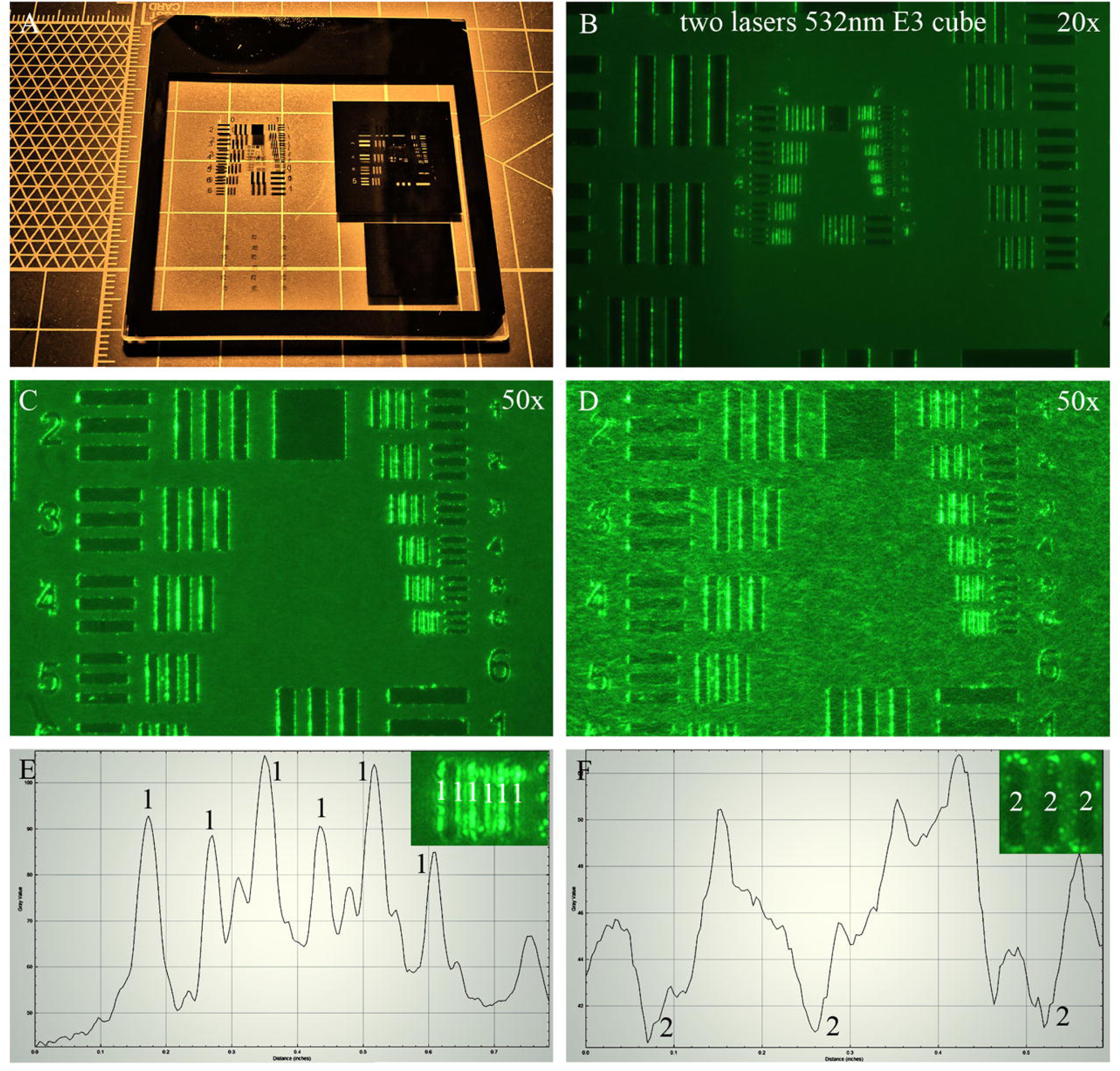
A - Overview image of a USAF 1951 slide used for measuring optical resolution. B - Image of the USAF 1951 test target using an LMPlan 20x objective; illumination was provided by 532 nm lasers (Leitz E3 filter cube). C - Detailed image of the USAF 1951 test target using an LMPlan 50x objective. The lines of element 6 in group 7 are clearly visible, indicating an optical resolution of 2.2 µm. D - The same area, but with the fiber optic vibrator turned off. A significant increase in image pixelation (noisiness) is noticeable. E.F shows a pixel intensities of vertical and horizontal lines of Group 7 Element 6 (graphics made in ImageJ program). E - the peaks correspond to the bright, protruding line; three lines — six peaks (two ribs for one vertical line, photo top right). F - the dips correspond to a dark, deep line; three lines — three dips (photo top left is rotated on 90 degrees). First photo was taken by Ipad10. All other photos were taken with a Sony IMX178 sensor.

## 4. Discussion

### 4.1. Parallels with Confocal Microscopy and the Importance of Micromanipulators

Optical fibers have been used in optical microscopy for quite some time; a detailed review on this topic can be found in work from as early as 20 years ago (Flusberg et al., 2005). Optical fibers can even be used as focusable objectives, but ultimately their function boils down to one thing: transmitting light over a distance from the source. However, in the aforementioned review the word ‘micromanipulator’ is not mentioned once. Similarly, in another newer and very high-quality book publication (Sanderson J., 2019) the term ‘micromanipulator’ is also absent. The original Ellis concept likewise makes no mention of micromanipulators. This leaves only one question: why not? What prevents the use of a micromanipulator for precise optical fiber positioning? The answer is: nothing. Moreover, in a somewhat modified form, micromanipulators are already used in confocal laser scanning microscopy.

As an example, the very first Leica TCS SP series confocal microscope (Fig. 16.A) already featured an optical fiber pathway between the lasers, housed in a separate module, and the scanning head itself. In this model, light from the optical fiber is focused through a pinhole onto a dichroic mirror and then reflected onto a motor-controlled galvanometer scanner mirror. In this sense, there are not many differences between the modified Ellis concept presented in this article and the optical scheme already used in Leica’s confocal microscopy. In both cases, motors with controlled steps are involved in the optical path with a single goal: to precisely position the light beam from the optical fiber (=light source) on its way to the objective.

**Figure 16.**
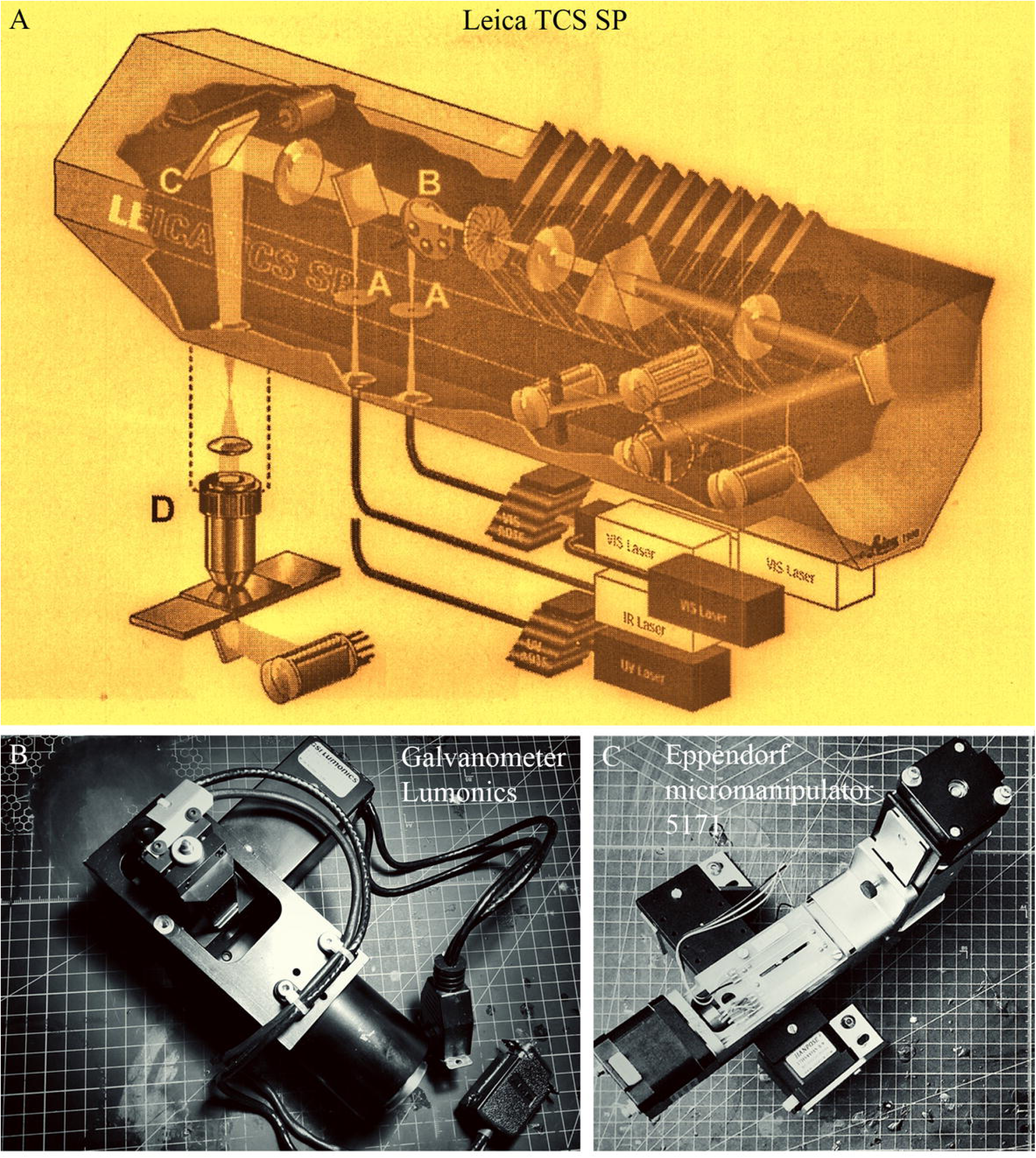
A - Schematic of the optical path in the Leica TCS SP laser scanning confocal microscope (image taken from Wright and Wright., 2002). The left part of the figure is of greatest interest, showing the light path from the lasers, via optical fibers connected to the confocal module housing. Within the module housing, the light beam passes through a pinhole (label A; its diameter can be comparable to the fiber optic core diameter) and is projected onto a dichroic mirror (label B), from which it is reflected to a galvanometer scanner mirror (label C). From this mirror, the beam is then reflected toward the objective (label D). B - Photo of the galvanometer scanner (Lumonics) from the Leica TCS SP confocal module. The mirror is housed within a sealed enclosure and is not visible, but it is controlled by two programmable motors, whose steps are precisely controlled via a driver board. C - Photo of the mechanical part of the Eppendorf 5171 micromanipulator with NEMA 17 stepper motors. Both the galvanometer scanner and the micromanipulator are components of the optical path that contain high-precision motors and are positioned directly before the objective in both cases, albeit on different sides of it depending on the type of optical system. The confocal microscope, with its lasers, galvanometer scanner, and optical fiber, shares many similarities in its opto-fiber-based components with the fiber-optic microscope; in places, the architecture is almost identical.

The galvanoscanner of a confocal microscope is a complete analog of a micromanipulator; they have similar functionality and a similar design (movable motorized axes perpendicular to each other, with a payload attached; Fig. 16.B). Essentially, the galvanoscanner mirror can simply be remounted onto a micromanipulator, for example, an Eppendorf 5171 (Fig. 16.C), and it can technically be positioned precisely using the manipulator in the same location as in the galvanoscanner, but with a caveat – the micromanipulator provides a degree of freedom along the axes that is unavailable to the galvanoscanner (three axes instead of two and increased linear travel of the axes instead of only 360-degree rotation of the mirror), especially when it comes to electronic micromanipulators like Eppendorf. It is precisely this freedom that allows the micromanipulator to be used in an unusual role – positioning an optical fiber in immediate proximity to the object, with minimal light loss – i.e., something that is inaccessible to the galvanoscanner due to its (albeit similar) design. Such use of manipulators has never been described before, where the manipulator (like the galvanoscanner) becomes an active participant in the microscope’s optical path. This is the novelty of the technology. And it is based on the Ellis concept, but with some differences.

### 4.2. Differences from the classic Ellis scheme

The concept of fiber optic lighting was slightly different in the already mentioned work of Ellis (1979). Firstly, this paper used single-mode fiber instead of multi-mode fiber. This proved to be more convenient, which was deduced experimentally. Multimode fiber required much higher motor speed frequencies (minimum 150Hz), which resulted in excessive vibration of the microscope. At lower frequencies using multimode fiber the speckle was maintained. Even physically separating the vibration unit from the microscope on different tables did not help. At frequencies of 10-20Hz in the case of single-mode fiber, the vibrations transmitted to the microscope were completely imperceptible.

The Ellis work uses a piezo element as a vibrator (100kHz), whereas in this work it is a vibration module from a 3D slicer on a stepper motor. The replacement of the piezo module with a mechanical module is due to the simplicity of the design decisions. Even the mechanism of an electric toothbrush can be used as a vibrator with a single-mode fiber, which was also experimentally tested (one of the B3-PPC samples was filmed in this mode, it is presented on the photo).

Finally, the presence of micromanipulators particularly distinguishes the concept of a fiber-optic fluorescence microscope from the original Ellis concept, where the tip of the optical fiber was on the zoom lens to the microscope, rather than directly on the object of study. A similar idea was implemented in Nightsea microscopes, but there the positioning of the optical fiber was carried out by means of an elastic harness.

Thus, the concept of fiber-optic microscope presented here is original and quite self-sufficient, we can say that it is a parallel version of the original concept from Ellis.

### 4.3. Comparison of classical fluorescence microscope concepts with a fiber-optic microscope. Freedom of component choice and impact on final cost

Thus, it can be said that the fiber-optic fluorescence microscope lies midway between classical fluorescence and confocal microscopy. From confocal microscopy, the modified Ellis concept takes lasers, fiber optics, and (partially) a step-motor-controlled fiber positioning system (a micromanipulator as an extended equivalent of a galvanoscanner). From fluorescence microscopy, the modified Ellis concept does not so much take as it rejects, yet ultimately obtains an identical result. Let’s list which parts of the classical fluorescence microscope were successfully replaced:

1. The high-voltage power supply unit for the mercury lamp with its ignition system is replaced by a significantly cheaper power supply for household appliances, available in any electronics supermarket, when it comes to powering inexpensive, low-power diode-pumped solid-state lasers. Although, for example, Helium-Neon lasers certainly require high-voltage power supplies, the choice of lasers remains with the laboratory. Therefore, the price range for this component can be very wide.
2. The mercury lamp with a fairly limited lifespan (100-200 hours) is replaced by a set of lasers with a similar spectrum. The choice of lasers is at the discretion of the laboratory. In this work, inexpensive diode-pumped solid-state lasers or laser pointers (all costing no more than 100$ each) were used, and expensive Helium-Neon or fiber lasers costing tens of thousands of dollars were not employed (it is important not to confuse a fiber laser with a laser with an attached optical fiber; a fiber laser uses optical fiber as its active medium). The problem with expensive lasers is that they are often unavailable for sale to private individuals and even organizations (in the Russian Federation). However, their use would certainly be an interesting continuation of this work. All this, in the end, also contributes to a significant price spread.
3. The collimator and Lamphouse with its internal system of focusing lenses also become unnecessary due to the replacement of the mercury lamp with lasers. However, a clarification is needed here. The diode-pumped solid-state lasers used in this work (price 50-100$) do contain an element such as a collimator, but it is almost always already part of the laser’s design and its price. Moreover, the collimator of a solid-state laser cannot be compared with the collimator of a fluorescence illuminator (epi-block) due to completely different sizes, types of materials used, and number of components (the price of a separately sold collimator for diode lasers is in the range of 10-20$), probably because it is made of plastic rather than glass, but it nevertheless performs its function.
4. Filter cubes can be used, or one can do without them, leaving only emission filters. In this work, when evaluating different markers, both possibilities were demonstrated (DiI was evaluated using filter cubes, DiD and B3-PPC using only an emission filter). The choice remains with the laboratory. Of course, the price of an emission filter is usually several times lower than that of a filter cube, as the savings come from the absence of an expensive dichroic mirror, excitation filter, and housing.
5. Optical fiber positioning system – micromanipulators. A classical fluorescence microscope does not have such an instrument; with some (very significant) allowance, the positioning screws for the mercury lamp in the Lamphouse could remotely be considered an analogue (since they also position the light source), but they have completely different accuracy (they are metric screws). In this work, in the case of the Leitz Laborlux microscope, inexpensive manipulators assembled from macrorails along the y-axis and SEMX-60AC optomechanical platforms along the x and z axes were used. The pitch of the macrorails is 1 mm per division of the gear scale, and the pitch along the x and y axes is 10 μm per division of the screw drive scale. Overall, this is sufficient for routine tasks. To improve accuracy, the macrorails along the y-axis can be replaced with another SEMX-60AC optomechanical platform. However, if there are no budget constraints, electronic micromanipulators from Sutter Instrument or Eppendorf are preferable. Using a frame made from aluminum metal construction kits, the heavy micromanipulator can be installed on the microscope body without much difficulty. The accuracy of the stepper motors in the micromanipulator (usually a Nema 17) is 400 steps per full shaft revolution. The micromanipulator’s electronics allow achieving an accuracy of 160 nm per one motor shaft step (this value was taken from the manual of the Eppendorf 5171 Micromanipulator). In the future, modifying the micromanipulator by using stepper motors with planetary gearboxes could further increase the accuracy. In any case, achieving exceptional precision in fiber optic positioning is not a difficult problem; all that is needed is to replace the stepper motors and install planetary gearboxes at the researcher’s discretion. For this reason, the price range for micromanipulators also remains at the laboratory’s choice, as it involves amounts from 100$ to several thousand dollars for a new Eppendorf set. Although, of course, the price for electronic manipulators can be significantly reduced if they are developed from scratch (this is a topic for a separate article) or if used models from eBay are utilized.
6. Vibration module – has no analogue in a fluorescence microscope, but overall it is a very inexpensive component, consisting of a motor and a linear translator, all of which typically falls within the 100$ range.

### 4.4. Versatile placement of the fiber optic fluorescent lighting system

The main array of the fiber optic unit consists of an aluminium metal constructor, The metal constructor is essentially a framework, or scaffolding, on which anything can be mounted, in this particular case fiber optic micromanipulators. The advantage of the aluminium metalconstructor is its versatility. The supporting body of any micromanipulator always has one or another technological holes in the casing. Most often their role is to fix the outer skin of the microscope. These technological holes can be used to attach the aluminium profile via corner connectors. The main thing is that the aluminium profile must be fixed firmly and contain a sufficient number of elements to support the weight of the manipulators. The number of parts and the layout of the profile are selected depending on the microscope model, and for this reason it will always be a slightly individual arrangement. The advantage is that the age of the microscope model or its type are not limiting factors. The Leitz straight and inverted microscope models produced more than 30 years ago were used in this work, and they proved to be absolutely suitable for modification. There is no reason why the same metalwork cannot be fitted to any other models.

Once the metal construction is installed, it is easy to install manipulators. And it can be not only manipulators based on optomechanical platforms, but also commercial ones, such as Narishige, Eppebdorf or manipulators from Sutter instrument. The aluminium profile makes it possible to install any set of manipulators from any company.

Finally, not needing a ploemopak module with filter cubes allows the optical system of the microscope to be greatly simplified. Usually, any top-four microscope (Nikon, Olympus, Leica, Zeiss) has slots for polarising or ND filters, and it is very easy to find a suitably sized interference extinction filter for them. Even if such a slot is suddenly not available, the interference filter can always be placed in the eyepiece adapter for the microscope camera.

All this combination of properties allows any researcher without special engineering skills to easily turn any microscope of transmitted light into a fluorescent microscope. And for this purpose nothing is required, except the correct arrangement of simple parts of the aluminium metal constructor, on which you can already attach any manipulators, both homemade and factory-made.

### 4.5. Limitations inherent in the fiber-optic fluorescence microscope

Like any instrument, a microscope has certain limits on the parameters within which it can be used. For a fluorescence microscope, there are two such parameters - spectrum width and magnification range.

1. Spectrum width. The visible spectrum of light ranges from 380nm to 780nm. In addition to the 488nm (blue laser), 532nm (green laser) and 650nm (red laser) laser pointers used in this work, there are also 405nm (violet laser), 450nm (blue-violet laser), 505nm (turquoise laser), 593nm (yellow laser) and 800nm (infrared laser) laser. Thus, there is no limitation on the spectrum width (due to laser pointers of different wavelengths) of the fiber-optic microscope, which was demonstrated by a line of corresponding lipophilic tracers.
2. Range of magnifications. In this work, the main array of photographs was taken at low magnifications of 2.5x-20x, which was due to the specificity of the study objectives (assessment of macro-distribution of labelled neuronal structures on the slice). The minor part of neuron photographs was taken at 50x magnification in order to demonstrate the capabilities of the new lighting system. One feature all these photos had in common was that they were all taken using lenses with a long working distance (9 to 12 mm). It was the large working distance of the lens (at least 5 mm) that allowed precise positioning of the fiber optic from a lateral position. For this reason (but not only), the use of low magnification and short working distance lenses is impractical, as is the use of 100x lenses, which almost always have a working distance of 2 mm. The exception here are the oil-free Mitutoyo lenses, for which even in the case of 100x magnification the working distance is 3-6 mm. But Mitutoyo lenses can not be mounted on every microscope.
3. Resolution limitations. The same as for a classical fluorescence microscope. This is confirmed by the ISAF 1951 test and the ability to detect individual 200 nm fluorescent beads.
4. The SDNR and CNR values for the different illumination systems are similar, with a slight advantage for laser illumination.
5. Field of view limitations – using two lasers results on average in a field of view approximately half the area of a classical fluorescence microscope (46.94% at 4x magnification). However, this ratio depends on both the magnification level (higher magnification reduces the difference in area) and the number of lasers (and optical fibers). Furthermore, as measurements with ФМЭЛ-1A show, fluorescent signal is still detected even at a distance of 140 µm from the visible edge of the illuminated area. The system allows for scaling up the number of optical fibers and lasers to increase the illuminated area within the visible field. The choice to implement this capability remains with the laboratory.
6. Variation in fluorescence quantum yield/brightness depending on objective magnification – an inevitable consequence of having the optical fiber positioned externally to the objective. At low magnifications (4x), the quantum yield under fiber illumination is higher than under epi-illumination via the fluorescence ploemopak block; at 10x magnification the values equalize; at 20x and below, epi-illumination via the fluorescence illuminator block yields a higher quantum yield. This is likely related to the epi-characteristics of the objectives. Objectives with low magnification and long working distances are apparently designed from the outset to capture the maximum number of photons from the illuminated area. In the case of epi-illumination via the objective, the light beam per unit area is probably more concentrated (as the objective acts as a condenser), whereas the fiber-optic microscope currently lacks such a concentrating mechanism at this stage (for now). Therefore, the quantum yield at higher magnifications will be lower.

### 4.6. Comparison of a Fiber-Optic Fluorescence Microscope with Smartphone-Based (SFM) Models

In addition to the classical fluorescence microscope, which has been the primary subject of comparison in this work, there is another modern type of fluorescence microscopy based on using smartphones as digital cameras, laser diodes, and specialized lenses, all housed within a 3D-printed enclosure (Smartphone-Based Fluorescence Microscope – SFM). As an example, we will specifically examine the model described in Sami et al., (2023). This comparison is meaningful in the context of cost and the competitive position of a fiber-optic system in the modern landscape. However, considering the points made above, the modularity of the fiber-optic system inherently implies an exceptionally wide price range for its components, especially regarding manipulators and lasers, whose cost can differ by orders of magnitude depending on a laboratory’s budget. Therefore, it is more rational to focus not so much on a price comparison, but rather on the technical characteristics and, more precisely, the limitations inherent to SFM systems, to draw relevant conclusions.

1. Lens instead of an Objective. The design of SFM systems does not accommodate the installation of multi-component, complex, and expensive objectives that provide maximum depth of field and sharpness. Can a smartphone lens be equated to a Zeiss Neofluar objective (costing ∼5000$ for each)? Furthermore, an SFM system apparently requires manual lens replacement (with complete disassembly of the housing) each time a magnification change is needed, as a turret mechanism is not provided.
2. Fixed Z-axis Positioning. The aforementioned model (Sami et al., 2023) has only 4 fixed Z-axis positions. What if a focus shift of a few microns is required for sample evaluation? The design of the SFM microscope excludes the possibility of fine Z-axis adjustment (a feature handled by a built-in, complex planetary reducer in a standard factory microscope).
3. Lack of a Mechanical Stage. The absence of a stage in SFM systems eliminates the possibility of moving the sample along the X and Y axes and evaluating large or non-standard sized objects. Evaluation seems possible only in the center of the field of view.
4. Unresolved Coherence Issues. It is unclear how using a laser diode solves the problem of coherence. The vibrating optical fiber in our system is needed precisely for this reason (to scatter highly coherent laser radiation). The absence of vibration results in a pixelated image with very high levels of noise (see results for the 1951 USAF test). How the issue of coherence reduction is addressed in the SFM system remains unclear.
5. Analysis Time Limitations. If the laser diode is battery-powered, this imposes strict limitations on operational duration. The fiber-optic microscope is much less dependent on power sources in this regard, although batteries can also be used in it if lasers are replaced with laser pointers.
6. Inflexibility in Wavelength Switching. It is unclear how, within a single SFM microscope housing, one can change the laser diode and emission filter when a different wavelength is needed. Is the concept implemented as: one housing – one diode?
7. Precision and Tolerance Limitations of 3D Printing. A fiber-optic microscope is always an add-on mounted on top; its main body is a pre-existing standard transmitted light microscope with all the necessary characteristics regarding part precision and tolerances inherent to a factory-made product. The lasers and high-precision manipulators are also factory-made items, and the aluminum frame (which integrates them into a single unit) is merely a supporting structure for these manufactured components. Can the precision of a plastic part printed on a 3D printer compare with that of a factory-assembled transmitted light microscope made of metal?

Conclusion: A fiber-optic microscope and an SFM microscope are systems with vastly different characteristics, sharing only one commonality – both deal with fluorescence. A direct comparison would be incorrect due to the excessive disparity in the range of their technical specifications.

## Summary

This work has presented a novel concept for a fluorescence microscope possessing unique characteristics, such as exceptionally precise positioning of the illuminated field and even the ability to divide this field into separate segments (thanks to micromanipulators). The most critical attributes of the entire system are its accessibility and universality (enabled by the aluminum frame structure). The fiber-optic microscope is in no way inferior to a classical fluorescence microscope, neither in resolution nor in the range of investigatable wavelengths, let alone to SFM models. The next logical step would be to explore the possibility of transitioning this concept into the realm of confocal microscopy (the capabilities for object scanning using micromanipulators naturally lead in this direction), but that is a topic for the next article.

## Author contribution

Klepukov A: author of the Assembled vibrating slicer and the article, the performer of the experimental part of the work on 99%. Sponsor of research on 100%. All work was made in DiI-lab (private company).

## Acknowledgments

The author of the work (Klepukov A) expresses gratitude to Lovat M.L (Scientific Research Institute of Mitineniring of the Moscow State University) for conducting the stage of perfusion and euthanasia of neonatal mice. The author of work expresses gratitude to Ivan Boldyrev and Elena Vodovozova for B3-PPC (Institute of bioorganic chemistry RAS). The author of work expresses gratitude to Tatyana Dragunova for the photo of the fiber-optic fluorescence microscope. The author of work expresses gratitude to Vadim Miroshkin for consulting about ФМЭЛ-1А.

The author of work also expresses gratitude to Julia Komarova for translating the whole text into English

